# Harnessing the central dogma for stringent multi-level control of gene expression

**DOI:** 10.1101/2020.07.04.187500

**Authors:** F. Veronica Greco, Amir Pandi, Tobias J. Erb, Claire S. Grierson, Thomas E. Gorochowski

**Author notes:** Correspondence should be addressed to T.E.G.

## Abstract

Strictly controlled inducible gene expression is crucial when engineering biological systems where even tiny amounts of a protein have a large impact on function or host cell viability. In these cases, leaky protein production must be avoided at all costs, but ideally without affecting the achievable range of expression. Here, we demonstrate how the central dogma offers a simple way to effectively address this challenge. By simultaneously regulating both transcription and translation, we show how relative basal expression of an inducible system can be greatly reduced, with minimal impact on the maximum induced expression rate. Using this approach, we create several stringent expression systems displaying >1000-fold change in their output after induction *in vivo* and up to a 350-fold change when used in a cell-free expression system. Furthermore, we find that multi-level regulation is able to suppress transcriptional noise and creates a digital-like switch when transitioning between ‘on’ and ‘off’ states. This work provides foundational knowledge and a genetic toolkit of parts to create multi-level gene expression controllers for those working with toxic genes or requiring precise regulation and propagation of cellular signals. It also demonstrates the value of exploring more complex and diverse regulatory designs for synthetic biology.

## Introduction

Since the development of the first inducible systems in the early 1980s ^1^, the ability to dynamically control gene expression through the use of small molecules ^2^, light ^3,4^, and other signals ^5^ has revolutionized biotechnology. From controlling shifts between cell growth and protein production stages during large-scale fermentations ^6^, to the detailed characterization of genetic parts and circuitry ^7^, the control of gene expression underpins a huge variety of applications. However, while switching expression of a gene ‘on’ or ‘off’ is conceptually simple, it is rare for genes to have such discrete states or ever be completely silenced. Stochastic effects ^8,9^ and leaky expression are widespread and potentially important for adaptation in natural systems but can wreak havoc in engineered systems where genes are toxic to a host or responses are highly sensitive and easily triggered by unavoidable fluctuations ^10,11^.

Early systems for controlling gene expression relied on repurposing native regulatory components such as transcription factors. One of the most widely used is the P_*tac*_ system ^1^. This consists of a constitutively expressed LacI repressor that can form dimers and tetramers to strongly bind operator sites within a P_*tac*_ promoter sequence and sterically block initiation of RNA polymerase (RNAP). LacI is sensitive to Isopropyl β-d-1-thiogalactopyranoside (IPTG) and at high concentrations, the DNA binding activity of LacI is abolished. This lifts repression of P_*tac*_ and leads to strong transcription of genes regulated by this promoter. While in most cases such systems offer strong repression, because such regulatory systems focus on a single step during protein synthesis (i.e. transcription), they are vulnerable to fluctuations in regulator production and the stochastic nature of biochemical reactions during gene expression ^9^.

Over the past decade, synthetic biologists have developed more advanced methods to control gene expression. These include engineered regulators based on DNA binding proteins such as zinc fingers ^12^, TALENs ^13^ and CRISPRi ^14^, RNA-RNA interactions ^15–17^, post-transcriptional/translational processes such as RNA and protein degradation ^18^, as well as using directed evolution to optimize existing inducible systems ^19^. This offers a wealth of options to more strictly regulate gene expression through the coupling of multiple forms of regulation (e.g. affecting both transcription and translation of a gene) to reduce unwanted expression and improve the robustness of a system to component failure. However, few examples of such multi-level regulation have been implemented to date ^20,21^. This has resulted in an unclear picture of how best stringent multi-level control can be achieved and the trade-offs that exist between performance, regulatory complexity, and cellular burden when designing these systems.

Here, we address this problem by systematically studying the combined use of transcriptional and translational regulators to stringently control protein expression. Using a combination of mathematical modelling and combinatorial genetic assembly, we are able to design, build and test a variety of synthetic multi-level controllers (MLCs) and elucidate the relative performance of each. These controllers all implement a coherent type 1 feed-forward loop (C1-FFL) regulatory motif (**Figure 1A**) that is commonly found in natural genetic systems and is known to enable more stringent control of an output but is rarely used when designing new expression systems ^22^. We show how MLCs offer advantages for many applications spanning the stringent control of protein expression to the accurate propagation of information in a cell ^23,24^ and demonstrate how applying modern synthetic biology tools to even simple regulatory systems can offer paths towards the precise and reliable control of biological systems.

**Figure 1:**
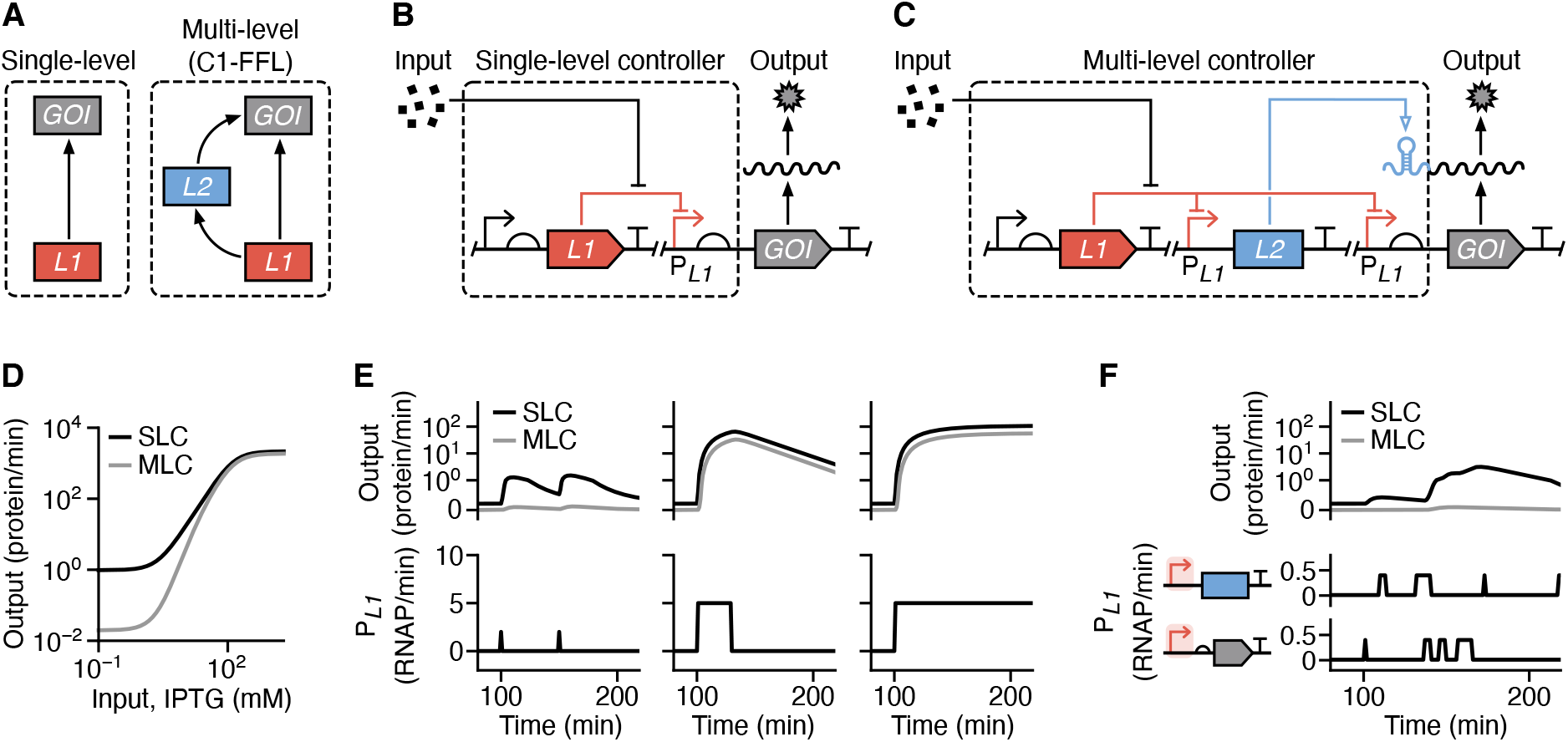
Stringent control of protein expression through multi-level gene regulation. (**A**) Two possible regulatory schemes to control the expression of a gene of interest (GOI): 1. control using a single regulator (*L1*), and 2. multi-level control using two separate regulators (*L1* and *L2*) connected in the form of a coherent type 1 feed-forward loop (C1-FFL). (**B**) Schematic of a genetic implementation of a single-level controller (SLC) that uses only transcriptional (red lines) regulation. An input (e.g. small molecule) modulates activity of the P_*L1*_ promoter and production of the GOI. (**C**) Schematic of a genetic implementation of a multi-level controller (MLC) that uses both transcriptional (red lines) and translational (blue line) regulation. An input (e.g. small molecule) modulates activity of the two P_*L1*_ promoters and an internal *L2* regulator activates the translation of GOI transcripts to finally produce the output protein. (**D**) Steady state response functions from mathematical models of the SLC and MLC. (**E**) Dynamic model simulations of the SLC and MLC and their response to different forms of temporal input (left to right): delta functions (P_*L1*_ activity = 2 RNAP/min for 1 min at 100 min and 150 min), a pulse (P_*L1*_ activity = 5 RNAP/min from 100–130 min), and a step function (P_*L1*_ activity = 5 RNAP/min from 100 min onwards). The activity of both P_*L1*_ promoters in the MLC is considered identical. (**F**) Dynamic model simulations of the SLC and MLC showing suppression of intrinsic promoter noise by the MLC. The two identical P_*L1*_ promoters for the *L2* regulator and GOI are separately driven by independent and biologically realistic bursty transcriptional activity profiles (**Methods**).

## Results

### Stringent control of gene expression by harnessing the central dogma

In most synthetic genetic circuits, control of gene expression is achieved through the use of a single type of regulation (**Figure 1A**), with control of transcription predominantly used. While this type of single-level controller (SLC; **Figure 1B**) is often sufficient for many applications, the central dogma naturally lends itself to more stringent multi-level regulation where both transcription and translation are controlled simultaneously (e.g. via transcription factors and RNA-based translational switches). Such multi-level controllers (MLCs; **Figure 1C**) can be generalised by a genetic design that consists of an *L1* gene encoding a level 1 transcriptional regulator with cognate promoter P_*L1*_, and an *L2* gene encoding a level 2 translational regulator. Both *L2* and the gene of interest (GOI) are separately transcribed by P_*L1*_ promoters and the product of *L2* activates translation of the GOI transcript. This MLC encapsulates a coherent type 1 feed forward loop (C1-FFL) in which both *L1* and *L2* are necessary for production of the GOI.

To explore the possible benefits of this regulatory motif, we developed mathematical models to capture how the rate of production of a GOI varied in response to differing concentrations of an input inducer for both the SLC and MLC designs (**Supplementary Note 1**; **Supplementary Data 1**). We generated steady state response functions by simulating the models using biologically realistic parameters (**Supplementary Table 1**) over a range of different input IPTG concentrations. As expected, the output production rate displayed a sigmoidal shape with both controllers reaching near identical maximum rates at high input IPTG concentrations (**Figure 1D**). The main difference was that the MLC design displayed a 50-fold lower output than the direct controller at low IPTG concentrations, leading to significantly reduced basal expression when no input was present (**Figure 1D**). This caused the MLC design to have both an increased dynamic range and fold-change between ‘off’ and ‘on’ states when compared to the SLC design.

**Table 1:**
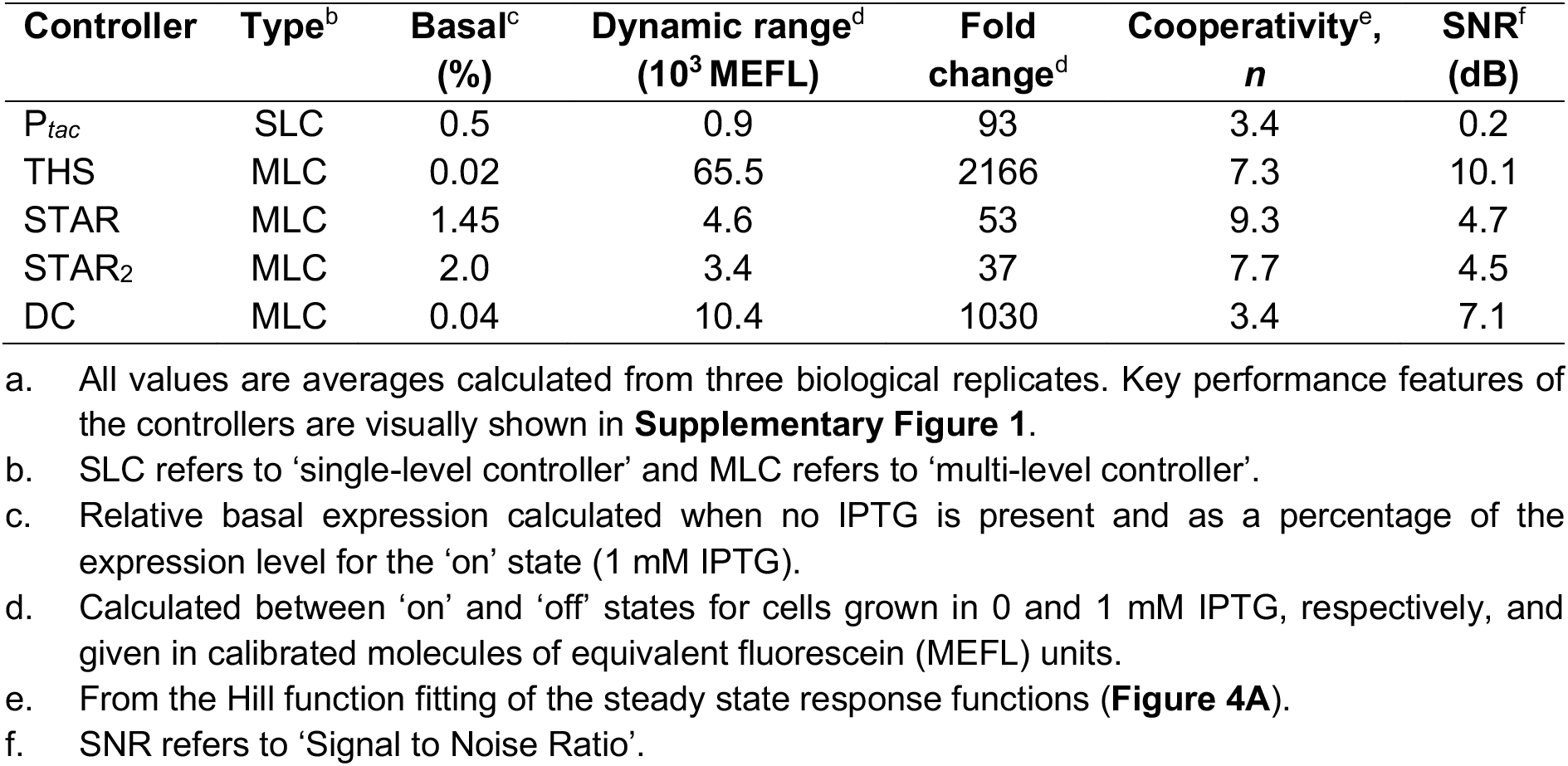
Performance summary of the single- and multi-level controllers *in vivo*^a^.

| Controller | Type<sup>b</sup> | Basal<sup>c</sup><br>(%) | Dynamic range<sup>d</sup><br>(10<sup>3</sup> MEFL) | Fold<br>change<sup>d</sup> | Cooperativity<sup>e</sup>,<br>*n* | SNR<sup>f</sup><br>(dB) |
| --- | --- | --- | --- | --- | --- | --- |
| *P<sub>tac</sub>* | SLC | 0.5 | 0.9 | 93 | 3.4 | 0.2 |
| THS | MLC | 0.02 | 65.5 | 2166 | 7.3 | 10.1 |
| STAR | MLC | 1.45 | 4.6 | 53 | 9.3 | 4.7 |
| STAR<sub>2</sub> | MLC | 2.0 | 3.4 | 37 | 7.7 | 4.5 |
| DC | MLC | 0.04 | 10.4 | 1030 | 3.4 | 7.1 |

We also simulated the output protein production rate for both models when exposed to a range of dynamic inputs. These included delta functions, as well as pulse and step inputs (**Figure 1E**). Simulations showed that both types of controller displayed virtually identical output responses for both the pulse and step inputs, with only a small reduction in output expression rate for the MLC that matched its lower basal expression level. However, significant differences were observed in the responses to the delta function input. While the SLC led to moderate sized pulses in output, the MLC design fully suppressed all output activity with only tiny fluctuations in the output expression rate observed. The behaviour of the MLC arose from the need for both *L1* and *L2* to be expressed to sufficiently high levels for expression of the GOI to be triggered. The short pulses of expression caused by the delta function input were insufficient to cause this switch and allowed the MLC to effectively filter out these transient events in its input.

The ability to filter out rapid fluctuations is particularly important for stringent control in systems where input promoters exhibit high levels of intrinsic noise. In such scenarios, protein levels can vary significantly across a population of cells ^8^ due to the often bursty nature of gene transcription. This is commonly seen for weak promoters where intrinsic noise dominates. Rather than the activity of a weak promoter being uniformly low, it instead displays short bursts of strong activity separated by long periods of inactivity ^8,25^. Across a population this averages out to a low overall expression level, but large variability is present between cells. As seen for the delta function inputs, such input profiles driving the SLC will lead to large fluctuations in the output. However, because intrinsic promoter noise is specific to an individual promoter and uncorrelated between multiple identical versions of a promoter within a construct, the MLC design which contains two copies of the input promoter P_*L1*_ should find that a burst of expression from one P_*L1*_ promoter is highly unlikely to occur at the same time as a burst from the other. Therefore, the MLC will suppress noise in the output.

To test this hypothesis, we generated accurate time-series promoter activity profiles based on a two-state model ^25^ where the mean length of time a promoter was in an ‘on’ active and ‘off’ silent state (Δ*t*_ON_ and Δ*t*_OFF_, respectively) were ⟨Δ*t*_ON_⟩ = 6 min and ⟨Δ*t*_OFF_⟩ = 37 min. These values were taken from previous experimental measurements in *E. coli* ^25^. We also set the activity of the P_*L1*_ promoter when in an ‘on’ state to a biologically realistic 0.25 RNAP/min. Independent time-series were generated for each P_*L1*_ promoter in the MLC and only one of these was used for the SLC where only a single P_*L1*_ promoter is present. These profiles were then fed into our existing dynamic models and the responses of the systems simulated. We found that the output production rate for the SLC saw large increases, especially where the input consisted of longer bursts of activity or several bursts in short succession (**Figure 1F**). In comparison, the MLC fully suppressed all output production making it an excellent filter of intrinsic promoter noise.

### A genetic template to explore multi-level gene regulation

There are many ways that an MLC could be implemented biologically. Furthermore, when implementing such a controller it is often necessary to switch the input that is used and internal regulators such that multiple controllers can be used simultaneously within the same cell. To meet these requirements, we developed an 8-part genetic template and toolkit of parts to allow for the rapid combinatorial assembly of MLCs (**Figure 2A**). The design enables both single and multi-level regulation, has the option to introduce protein tags for further post-translational control of the GOI (e.g. through signalled degradation) and is structured to minimise the chance for transcriptional readthrough to cause unwanted expression of the component parts. The toolkit comprises 8 types of part plasmid (pA–pH) and a backbone plasmid (pMLC-BB1) in which the final MLC design is inserted (**Supplementary Data 2**). Assembly is performed using a standard one-pot Golden Gate reaction with individual blocks designed to use 4 bp overhangs with minimal cross reactivity to ensure the correct and efficient ligation of parts ^26^. Furthermore, rapid screening of successful inserts is enabled by the drop-out of an orange fluorescent protein (*ofp*) expression unit ^27^ (**Supplementary Note 2**).

**Figure 2:**
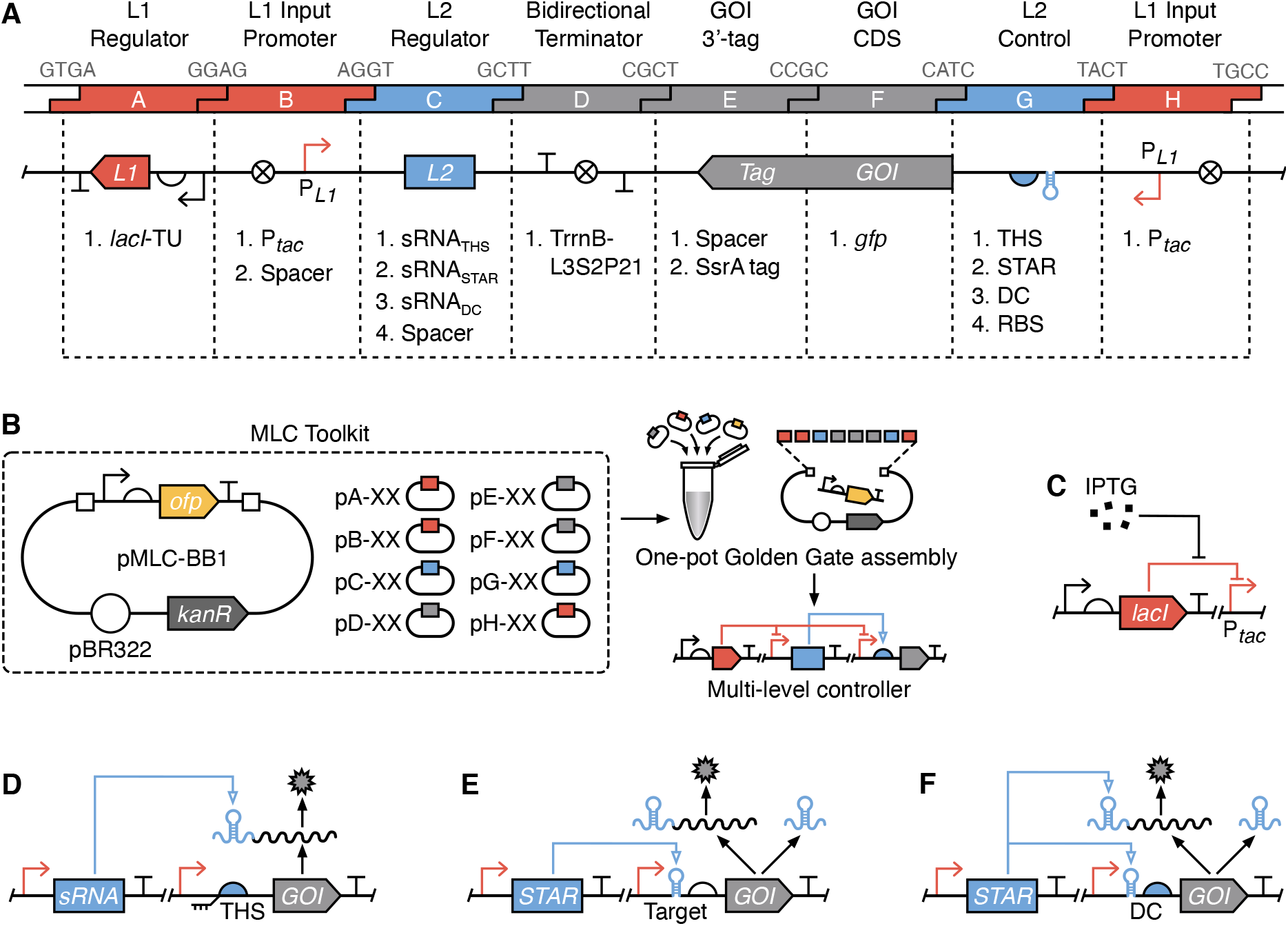
Combinatorial assembly of gene expression controllers. (**A**) Summary of the 8-part genetic template used to allow for systematic exploration of direct and multi-level gene regulation. The 4 bp overhangs used for Golden Gate assembly are shown in grey at their respective junctions. Available genetic elements are listed below each corresponding part type (A–H). (**B**) The MLC toolkit contains a set of plasmids that can be combined using Golden Gate assembly to create a variety of direct and multi-level controllers (**Supplementary Figure 3**). (**C**) The *lacI* transcription factor responsive to IPTG used for level 1 (*L1*) transcriptional regulatory control. (**D**) Toehold switch (THS) translational regulator used for level 2 (*L2*) control. (**E**) Small transcription activating RNA (STAR) transcriptional regulator used for *L2* control. (**F**) Dual control (DC) transcriptional and translational regulator used for *L2* control.

Using this toolkit, we aimed to compare the *in vivo* behaviours of different SLC and MLC designs with a focus on the different mechanisms that could be used for *L2* control and the affect these might have on overall performance. For the SLC design we chose the widely used P_*tac*_ system introduced earlier (**Figure 2C**). To simplify comparisons, we also used the P_*tac*_ system for *L1* control in all the MLC designs and combined it with three different RNA-based *L2* regulators. These included a toehold switch (THS; **Figure 2D**) ^17,28^, a small transcription activating RNA (STAR; **Figure 2E**) ^29^, and a dual control system (DC; **Figure 2F**) ^21^.

The THS regulator encodes a structural component followed by a ribosome binding site (RBS) that is used to drive translation of the GOI (**Figure 2D**). The structural region is designed to form a strong hairpin loop that when transcribed hinders the ability for ribosomes to bind the RBS, and thus inhibits translation. Translation is activated by expression of a complementary small RNA (sRNA) trigger that hybridizes to a short unstructured region of the THS which causes a breakdown in its secondary structure. This conformational change allows ribosomes to bind the RBS and translation of the GOI to proceed. THSs were selected because they offer strong repression of translation, can be designed computationally, and large libraries of designs exist with minimal crosstalk when used together ^17,28^.

Unlike the THS, the STAR regulator works at a transcriptional level. The STAR’s target is placed before an RBS in the 5’ untranslated region (UTR) of the GOI (**Figure 2E**). This forms an intrinsic terminator when transcribed and inhibits GOI expression. Activation is achieved by expression of the STAR RNA, which interacts with the target, prevents terminator formation and thus allows for expression of the downstream GOI. Similar to THSs, STARs have been shown to offer strong repression and there exist large libraries of orthogonal variants ^16,29^.

Finally, the DC regulator combines both transcriptional and translational control by modifying the pT181 attenuator ^21^. The DC target is placed in the 5’ UTR of the GOI and encodes an intrinsic terminator that includes the RBS (**Figure 2F**). When transcribed, the intrinsic terminator not only halts transcription, but also represses translation by causing the RBS to form a strong RNA secondary structure making it inaccessible to the ribosome. Activation is achieved by expression of a STAR, which interacts with the target, both preventing terminator formation and causing a conformational change in the RNA structure that makes the RBS accessible for translation initiation. The DC regulator was chosen due to this combined regulatory action which has been shown to produce strong repression ^21^. However, to date, only a single of these regulators has been created, limiting future applications.

DNA encoding parts for each of these regulatory systems was synthesised and our toolkit used to assemble the SLC and three MLC designs. Superfolder green fluorescent protein (*gfp*) was chosen as the GOI to allow for the measurement of output expression in single cells using flow cytometry.

### Performance comparison of the controllers

To characterise the performances of the controllers, we transformed *Escherichia coli* cells with each construct and measured GFP fluorescence using flow cytometry for ‘off’ and ‘on’ input states. As P_*tac*_ was used as an input for all the designs, this corresponded to growing the cells in either 0 or 1 mM IPTG, respectively (**Methods**). Data from these experiments was then used to calculate the dynamic range and fold change in output GFP fluorescence (**Table 1**; **Supplementary Figure 1**).

We found a clear separation between output states for all designs with little variation between biological replicates (**Figures 3A**). All MLCs (THS, STAR and DC) reached higher expression levels than the P_*tac*_ SLC, and the THS and DC designs achieved large >1000-fold changes between output states. Notably, while the STAR design reached a much higher ‘on’ state than the P_*tac*_ design, the STARs high levels of basal (leaky) expression when no input was present resulted in a 43% lower fold change (**Table 1**).

**Figure 3:**
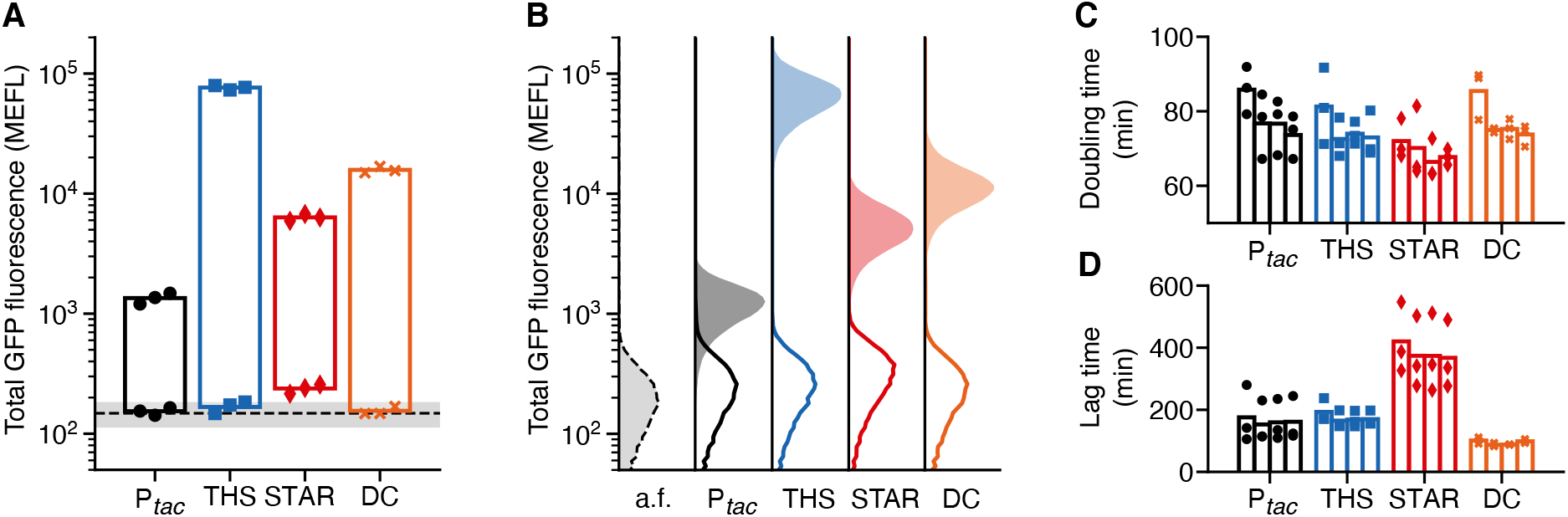
Performance comparison of single- and multi-level controllers *in vivo*. (**A**) Total GFP fluorescence for ‘off’ and ‘on’ input states (0 and 1 mM IPTG, respectively). Points show the three biological replicates for each controller and condition (black circles, P_*tac*_; blue squares, THS; red diamonds, STAR; orange crosses, DC). Black dashed line denotes the mean fluorescence of cell autofluorescence (a.f.) controls containing no plasmid with grey shaded region showing ± 1 standard deviation of 11 biological replicates. Fluorescence given in calibrated molecules of equivalent fluorescein (MEFL) units. (**B**) Flow cytometry distributions of total GFP fluorescence for ‘off’ (line) and ‘on’ (shaded) input states. Cell autofluorescence (a.f.) controls containing no controller are shown by black dashed line and light grey filled distributions. (**C**) Doubling time of cells harbouring direct and multi-level controllers for varying concentrations of IPTG (bars left to right for each design: 0, 0.1, 1, 10 mM IPTG). (**D**) Lag time calculated as the time to reach an OD_600_ = 0.15 after inoculation of cells harbouring controllers for varying concentrations of IPTG (bars left to right for each design: 0, 0.1, 1, 10 mM IPTG).

A challenge when calculating these measures (especially fold change) is the ability to accurately quantify very low level of output GFP fluorescence, which are near or identical to the autofluorescence of the cells. To better understand this aspect, we measured the GFP autofluorescence of untransformed *E. coli* cells, performing 11 biological replicates to estimate a fluorescence distribution that could be used as an approximate detection limit. Overlaying the average and standard deviation of the cell autofluorescence onto our results (**Figure 3A**, dashed line and grey shaded region), we found that the ‘off’ states for the P_*tac*_, THS and DC designs all fell within this region and very close to the average suggesting they have virtually no leaky expression at all.

Another difficulty when comparing the performance of the controllers is the need to consider the large differences in the maximum expression rates (e.g. >60-fold difference between the P_*tac*_ and THS designs for the ‘on’ state). It should be noted that the same P_*tac*_ promoter is used as input to all our designs and that it includes a 15 bp upstream spacer element to insulate its function from contextual effects arising from differing nearby sequences that are present in each design ^24,30^. It is therefore reasonable to expect the dynamic range of the input promoter’s transcriptional activity to be similar for each controller, with differences in output protein expression rate related directly to the different strength ribosome binding sites found in each *L2* regulator or the SLC design. Given these differences and to allow for an unbiased comparison, we calculated the relative basal GFP expression level of each controller as a percentage of its maximum output (**Table 1**). This showed that the THS performed best, displaying a 25-fold decrease in relative basal expression compared to the P_*tac*_ SLC with 0.02% relative basal expression compared to 0.5%, respectively. The DC design also performed well with 0.04% relative basal expression, while the STAR MLC saw the largest relative basal expression of 1.45%, nearly 3 times that of the P_*tac*_ SLC.

While comparisons of average expression levels between ‘on’ and ‘off’ states are useful, they are not able to capture the role of cell-to-cell variability inherent in all gene expression ^9^. Such variation is crucial when assessing the performance of stringent expression systems because even though average output states might be sufficiently separated to be distinguished, cell-to-cell variation across a population can lead to overlaps in the output distributions. Cells falling in this overlap are impossible to classify resulting in some cells with an undetermined state. Engineers have developed measures to help characterise the strength and quality of a signal (i.e. the ability to distinguish ‘on’ and ‘off’ output states) with the Signal to Noise Ratio (SNR) commonly used in other fields such as electronics. SNR has also recently been adapted for use when studying engineered genetic systems making it easier to understand how the quality of signals in a circuit are maintained or degraded as they pass through various genetic devices ^23^.

Using the flow cytometry distributions, we calculated the SNR for each controller in decibel (dB) units (**Table 1**; **Methods**). We found that the P_*tac*_ SLC performed worst with a low SNR of 0.2 dB, corresponding to a signal barely larger than the noise. This was evident for the flow cytometry distributions where a sizable overlap in the ‘on’ and ‘off’ states was seen (**Figure 3B**). All MLCs performed better with the THS achieving an SNR >10 dB. This improved performance was also evident from the flow cytometry data with clear gaps of varying sizes between the ‘on’ and ‘off’ output distributions (**Figure 3B**). This clearer separation between cells in an ‘on’ and ‘off’ state would make these parts ideal for genetic logic circuits, ensuring signals are cleanly propagated.

### Burden of controllers on the host cell

There is a growing awareness of the importance of considering the burden that engineered genetic parts and circuits place on their host cell ^31^. The introduction of a genetic construct that sequesters large quantities of shared cellular resources like ribosomes or heavily impacts core metabolic fluxes can lead to reduced growth rates and trigger stress responses that impair the function of engineered genetic parts ^32–37^. When designing the MLCs, we purposefully selected RNA-based regulators as previous results suggest that they impose a small metabolic burden on the cell ^38^. However, to experimentally verify this in our cells, we generated growth curves for all SLC and MLC designs (**Supplementary Figure 2**). Because the metabolic demands of the controllers would vary based on the concentration of inducer present (due to the varying levels of sRNA or STAR produced), cells were exposed to 4 different concentrations of IPTG (0, 0.1, 1, 10 mM) spanning the ‘off’ and ‘on’ states of the controllers.

From these growth curves, we estimated the doubling time during the exponential growth phase (**Methods**). We found that the SLC and all MLCs displayed similar doubling times of ~70 min (**Figure 3C**). Furthermore, we saw a slight decrease in the doubling times of all controllers as the IPTG concentration increased. This trend is counterintuitive given that an increasing IPTG concentrations will cause expression of the GOI and any *L2* regulators, increasing the burden on the cell. However, it is known that IPTG can have unexpected effects on cell physiology ^39^ and cause changes in plasmid stability ^40^, which could lead to reduced overall burden due to fewer copies of the controller plasmid or more efficient utilisation of available nutrients by the cell.

We also measured the lag time after inoculation into fresh media before the cells entered exponential growth (**Methods**). We found differences between many of the controllers with a lag time of ~165 min for the P_*tac*_ and THS designs, a shorter lag time of 88 min for the DC design, and a significantly longer lag time of 373 min for the STAR design (**Figure 3D**). Closer inspection of the growth curves showed that the DC design had a consistently higher initial cell density (optical density at 600 nm of 0.07 compared to 0.04 for the THS design), which could account for the shorter lag phase (**Supplementary Figure 2**). For the STAR design the elongated lag phase coincided with a consistently longer additional time of ~100 min to reach saturation of the culture.

To better understand if the extended lag phase of the STAR-based MLC was a general feature to be expected when using this type of regulator, we rebuilt this construct using a different STAR (STAR_2_) that had an identical initial 72 bp sequence, but unique 10 bp sequence at its 3’-end (**Supplementary Table 2**). As we would expect for such a similar design, testing of the STAR_2_ construct showed similar performance to the initial STAR design with a good dynamic range and similar leaky expression in its output (**Table 1**; **Supplementary Figure 3**; **Methods**). However, unlike the original, the STAR_2_ design displayed a lag phase (161 min) and doubling time (72 min) that closely matched the other MLCs. This suggests that long lag times observed for the original STAR design were likely due to some highly specific and uncharacterised off-target interactions with endogenous cellular processes and not due to a general feature of the STAR’s regulatory mechanism.

### Digital-like transitions and suppression of weak input signals

Our previous modelling of the MLCs showed that in addition to improved performance in ‘on’ and ‘off’ states, the addition of the *L2* regulator also altered the response function, causing a sharper transition from an ‘off’ to ‘on’ state due to the lower basal expression, and an ability to suppress low level noise in the input (**Figure 1D, E**).

To assess if these features were present, we generated response functions of the controllers by growing the cells in varying concentrations of input inducer and measuring steady state output GFP fluorescence. The sharpness of the transition is captured by the cooperativity of Hill function fits to this data. We found that in comparison to the P_*tac*_ SLC, both the THS and STAR MLCs saw more than a doubling in this value from 3.4 to more than 7, while the DC design maintained an identical value (**Table 1**). High cooperativities correspond to a very sharp step-like transition between ‘on’ and ‘off’ states that is clearly evident from the response function curves (**Figure 4A**). The high non-linearity in the response functions of the THS and STAR MLCs is potentially useful for information processing tasks. In particular, implementing digital logic within cells requires clear ‘on’ and ‘off’ states and limited chance for signals to reside at intermediate states. Sharp transitions in the response function ensure that there is less room for an input to fall at an intermediate point during the transition, ensuring an ‘on’ or ‘off’ state is always given. Furthermore, a high non-linearity can also be exploited to generate bimodality. For example, if a noisy input is positioned to span the transition point in the response function, a population of cells will have large groups of cells in ‘on’ and ‘off’ states, with much fewer in intermediate states because of the sharp transition and small probability of falling in this small region.

**Figure 4:**
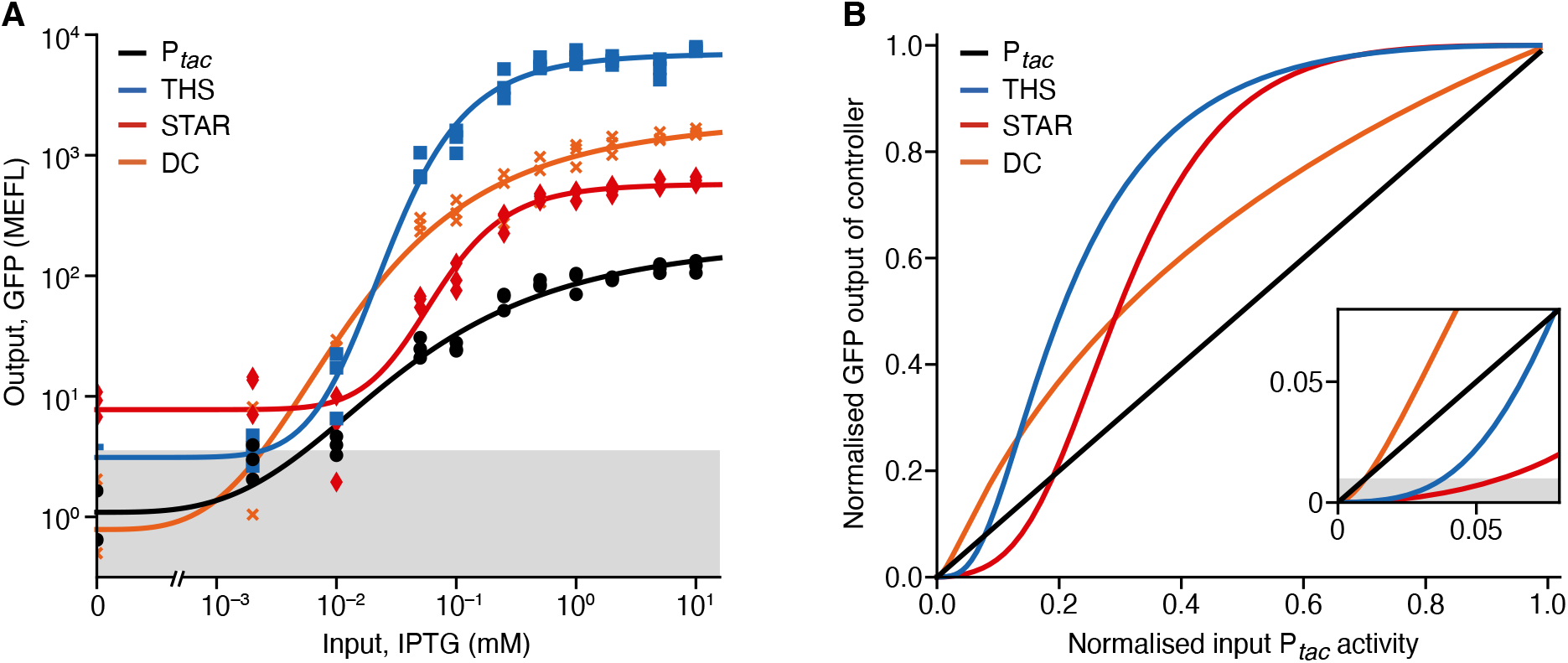
Response functions of single- and multi-level controllers *in vivo*. (**A**) Steady state response functions of the controllers showing output GFP fluorescence (corrected for cell autofluorescence) for varying input IPTG concentrations (0, 0.002, 0.01, 0.05, 0.1, 0.25, 0.5, 1, 2, 5, 10 mM). Points show the three biological replicates for each controller and condition (black circles, P_*tac*_; blue squares, THS; red diamonds, STAR; orange crosses, DC). Grey shaded region shows the standard deviation of cellular GFP autofluorescence from 11 biological replicates. (**B**) Comparison of how normalised GFP output (as a fraction of the maximum GFP fluorescence) varies in response to changes in the normalised transcriptional activity of P_*tac*_ (as a fraction of its maximum activity). Multi-level regulation can lead to the suppression or amplification of the output GFP production rate compared to direct transcriptional regulation (i.e. a specific multi-level controller’s line falls below or above the diagonal, respectively). Insert shows zoomed area and grey shaded region denotes a GFP output level of 1% for the controller.

To quantify the ability of each MLC to suppress low level input noise, we further analysed the response functions. As mentioned earlier, the large differences in dynamic range make comparisons between designs difficult. Given that the promoter driving transcription for each MLC is identical (P_*tac*_), the discrepancies arise from differing *gfp* translation rates controlled by the associated ribosome binding sites. These do differ in sequence and strength for each design and in some cases are specific and integral to the RNA regulator’s function. Therefore, to allow for comparisons, we normalised the output of each MLC to its maximum output and used data from the P_*tac*_ SLC to estimate the input activity of the P_*tac*_ promoter used in each controller. If no secondary regulation was present (as in the SLC), then we would expect the normalised input and output to follow a straight line where one equals the other (see P_*tac*_ design in **Figure 4B**). However, if the secondary regulation suppresses the input P_*tac*_ activity then a lower normalised output to input will be seen, and conversely, an amplification of the input will lead to a higher normalised output to input.

Using this approach, we assessed the responses of each MLC and found that all caused a suppression of low levels of input promoter activity and an amplification of higher input activities. This effect was most prominent for the THS and STAR designs, with both able to ensure controller output is maintained below 1% even when the input promoter reaches 3.5% activity (**Figure 4B**, insert). These results confirm the findings of our modelling and demonstrate the potential for using MLCs to filter out unwanted input activity in noisy environments.

### Controller performance in a cell-free expression system

There has been growing interest in the use of cell-free protein synthesis (CFPS) systems ^41^ as a means to prototype synthetic genetic circuits ^42^, enable the rapid characterisation of genetic parts and metabolic pathways ^43^, and more recently as a novel bioproduction platform^44^. While great progress has been made in expanding the applications of CFPS systems ^45–48^, strategies to stringently control protein expression have yet to be developed.

To assess the performance of our controllers in a cell-free context, we used a CFPS system created from crude *E. coli* cell lysate and performed simple batch reactions (**Methods**) that we continuously monitored so that output expression rate could be inferred from changes in GFP fluorescence over time. These experiments showed that all controllers were also functional in the CFPS system and behaved qualitatively similar to the *in vivo* situation (**Figure 5A**; **Supplementary Table 3**). Overall, MLC designs performed better than the SLC design, by showing a lower percentage of basal expression, a larger dynamic range and fold changes between ‘off’ and ‘on’ states with sharper, more digital-like, transitions (i.e. higher co-operativity in Hill function fits).

**Figure 5:**
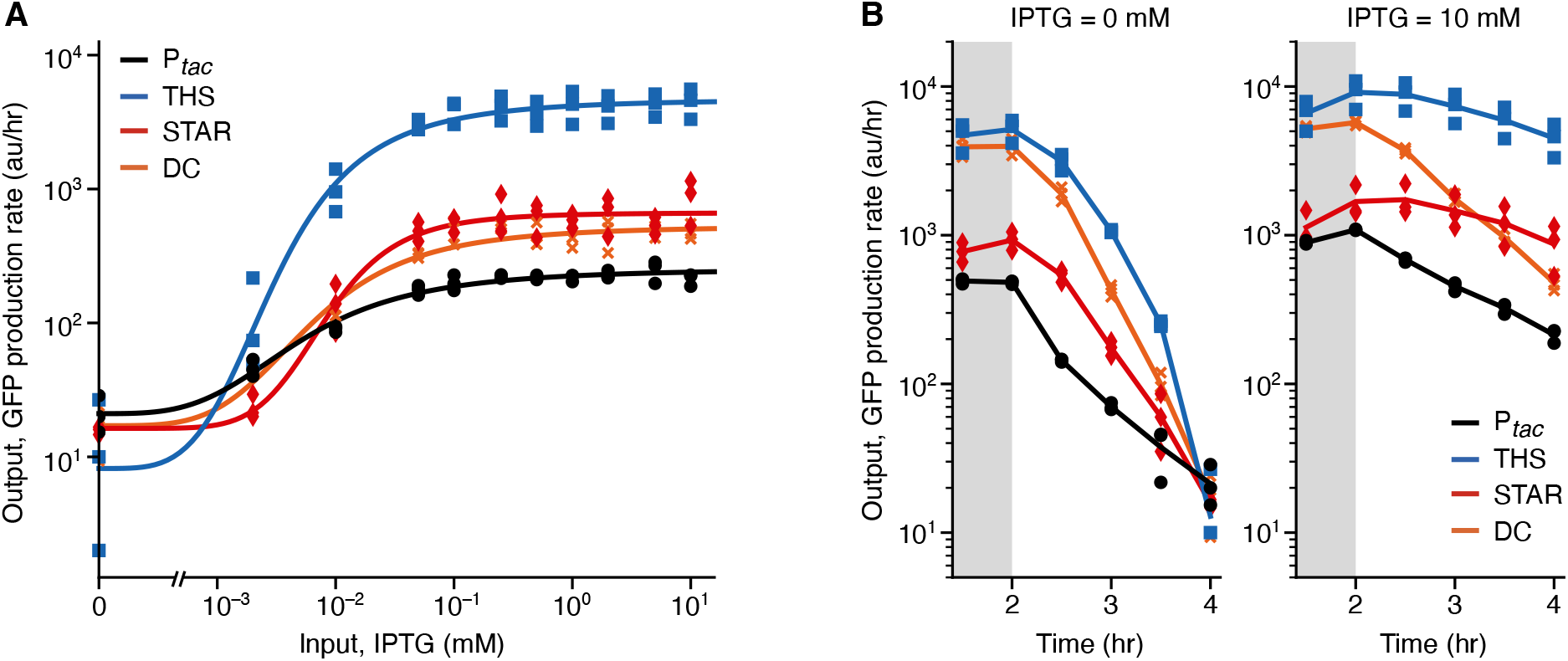
Performance of single- and multi-level controllers in a cell-free expression system. (**A**) Response functions of the controllers showing output GFP production rates in arbitrary fluorescence units per hour (au/hr) at 4 hours after the start of the cell-free reaction for varying input IPTG concentrations (0, 0.002, 0.01, 0.05, 0.1, 0.25, 0.5, 1, 2, 5, 10 mM). Points show the three biological replicates for each controller and condition (black circles, P_*tac*_; blue squares, THS; red diamonds, STAR; orange crosses, DC) (**B**) Output GFP production rate of the controllers over time since the start of the reaction. Time courses shown for controllers in an ‘off’ (0 mM IPTG; left) and ‘on’ (10 mM IPTG; right) state. GFP production rates at each time point calculated as an average GFP production rate over the previous 1.5 hours (**Methods**).

However, compared to the *in vivo* situation, we observed distinct differences in performance. The largest drop in performance was observed for the P_*tac*_ SLC design, for which basal expression reached 10% of the maximal output and only a 10-fold dynamic range (i.e. between ‘off’ and ‘on’ output states). Performance losses were also observed for the other MLC designs; however, the THS design displayed <1% basal expression and a ~350-fold change between ‘off’ and ‘on’ output states. These performance losses were likely caused by the relatively low effective concentrations of regulators that can be achieved in a CFPS system compared to the highly crowded cytoplasm of a living cell ^49^. Especially low concentrations of the LacI repressor will likely limit the maximal repression that can be achieved in the CFPS system, accounting for the higher basal expression observed in performance observed.

Notably, the STAR MLC showed a similar performance in respect to percentage of basal expression, fold-change and cooperativity compared to the *in vivo* setting (**Table 1** and **Supplementary Table S3**). This robustness may stem from the fact that the STAR design exploits secondary control at a transcriptional level, specifically, through premature termination of transcription in the 5’ UTR of the output gene, and therefore regulation limits the potentially active transcripts that are present within the reaction (**Figure 2E**). In contrast, the THS design produces full length transcripts and relies on continuous suppression of translation initiation by RNA secondary structures (**Figure 2D**). Our data suggests that the STAR regulator is less affected by the differing environment of the CFPS system than the THS, enabling the STAR design to maintain virtually identical performance across these contexts.

Time course measurements from these experiments also allowed us to quantify the output GFP production rate as the reaction proceeded. This data revealed a key difference between the *in vivo* and CFPS system that was observed for all controller designs. For the first 2 hours the expression rate for ‘off’ and ‘on’ states for each design were virtually identical, with regulation only being observed after this point and strongly affecting output GFP production rate after 4 hours (**Figure 5B**). The initial constant output GFP production rates in the CFPS system matched the order of different RBS strengths measured *in vivo*. The more rapid decrease in expression observed for the MLC designs versus the SLC for the regulated ‘off’ state is expected because of the additional regulatory layer (*L2* regulator) of these designs. The observed ‘lag’ phase of the regulation reflects very likely the time required for each controller to express sufficient LacI to interact with the P_*tac*_ promoters that act as the input in all our designs. In contrast to the CFPS system, in the *in vivo* experiments the cells had reached exponential growth and the systems were at steady state equal with LacI degradation and dilution rates equalling production rate to keep repressor concentration constant. Therefore, while multi-level regulation offers greatly improved control over gene expression in CFPS systems, for batch reactions, it is crucial that necessary regulatory components (e.g. repressor proteins) are present at sufficient concentrations from the start of an experiment to enable stringent regulation. This could be achieved by generating the CFPS system from cells that already express the regulators at high concentrations, by separately adding these components into the reaction mix before an experiment starts, or by making use of microreactors to enable the CFPS system to maintain steady state concentrations of regulators through continual dilution of the reaction products ^50^.

## Discussion

In this work we have shown how multi-level control of gene expression offers a means to more stringently regulate gene expression both *in vivo* and *in vitro*. By harnessing the multi-step process of transcription and translation that underpins the central dogma of biology and simultaneously regulating both processes in response to an input signal, we demonstrate through modelling (**Figure 1**) and experiments (**Figures 3**–**5**) how inducible expression systems can be created with greatly reduced leaky expression when in an ‘off’ state, while also maintaining high expression rates once induced. Furthermore, we have shown that multi-level regulation creates a more digital-like switch when transitioning between ‘off’ and ‘on’ states and suppresses low-level transcriptional noise (**Figure 4**), both of which are valuable properties when developing genetic systems for information processing or when highly toxic products or excitable systems act as downstream products.

Our top MLC design, which makes use of a THS for *L2* regulation, achieved >2000-fold change in output upon induction *in vivo* and displayed a 10 dB SNR (**Table 1**) making it one of the most tightly controlled and high-performance induction systems built to date. Furthermore, the flexibility of our modular genetic toolkit for assembling new multi-level controllers (**Figure 2**), and the availability of many other THSs, makes it easy to develop additional orthogonal MLCs that could be used in parallel within the same cell. It is worth noting that the underlying P_*tac*_ promoter that the THS MLC uses, achieved only a 93-fold change and 0.2 dB SNR when used alone as an SLC. Therefore, employing the multi-level regulatory approach outlined in this work could offer a means to greatly improve the performance of many existing low-performance transcriptional sensors, without any need to modify the transcription factors or promoter sequences making up these devices.

With the improvements we see when employing multi-level regulation, it is likely no coincidence that small interfering RNAs (siRNAs) are also widely used by bacteria to refine the regulation of many endogenous processes ^51–53^. RNAs are perfectly tailored for this task, imposing a small metabolic burden and offering a fast response. In this work, we selected synthetic RNA-based regulators that function through RNA-RNA hybridisation alone. While this reduces our dependencies on other cellular machinery and makes them easier to transfer between strains/organisms, it is known that many endogenous siRNA regulators employ protein chaperones such as Hfq to increase their binding affinity to targets and strengthen their regulatory effect ^54^. It would be interesting to explore the use of synthetic regulators that make use of these chaperones ^38^ or exploit recent advances in the RNA part design ^16^ to see whether further improvements in performance are possible.

The stringent regulation of our controllers is achieved by incorporating a C1-FFL regulatory motif that is known to be evolutionarily selected in many natural and engineered systems ^55^ and can be used to implement many useful functionalities ^22^. More recent work has also demonstrated the importance of interconnections and clustering of many motifs in coordinating more complex behaviours ^56,57^. While this work focused on demonstrating that transcriptional and translational regulation can fit neatly into a C1-FFL structure, an intriguing future direction would be to explore how these higher-level structures (e.g. motif clusters or higher-level network structures) might be implemented using the approaches outlined in this work to aid the coordination of multiple interrelated processes in parallel.

This study started with the goal of more stringently controlling gene expression. However, through the design of our MLCs it became evident that the more intricate regulatory designs we built had many other benefits. Synthetic biology to date has often focused on simplifying complexity and reducing systems to their minimal parts. Our findings indicate that complementary studies exploring the complexification of synthetic regulatory systems might also reap rewards allowing us to more efficiently exploit the capabilities of biology by combining many diverse processes and parts in unison. The genetic toolkit presented here offers a starting point for such studies focused on the fundamental processes of transcription and translation.

## Methods

### Strains, media and chemicals

All cloning and characterization of genetic constructs was performed using *Escherichia coli* strain DH10-β (Δ(ara-leu) 7697 araD139 fhuA ΔlacX74 galK16 galE15 e14-ϕ80dlacZΔM15 recA1 relA1 endA1 nupG rpsL (StrR) rph spoT1 Δ(mrr-hsdRMS-mcrBC) (New England Biolabs, C3019I). Cells were grown in DH10-β outgrowth medium (New England Biolabs, B9035S) for transformation, LB broth (Sigma-Aldrich, L3522) for general propagation, and M9 minimal media supplemented with glucose (6.78 g/L Na_2_HPO_4_, 3 g/L KH_2_PO_4_, 1 g/L NH_4_Cl, 0.5 g/L NaCl (Sigma-Aldrich, M6030), 0.34 g/L thiamine hydrochloride (Sigma T4625), 0.4% D-glucose (Sigma-Aldrich, G7528), 0.2% casamino acids (Acros, AC61204-5000), 2 mM MgSO_4_ (Acros, 213115000), and 0.1 mM CaCl_2_ (Sigma-Aldrich, C8106)) for characterization experiments. Antibiotic selection was performed using 100 μg/mL ampicillin (Sigma-Aldrich, A9518) and 50 μg/mL kanamycin (Sigma-Aldrich, K1637). Induction of the expression systems was performed using varying concentrations of isopropyl β-D-1-thiogalactopyranoside (IPTG) (Sigma-Aldrich, I6758).

### Assembly of controllers

All part plasmids were either directly synthesised (GeneArt, Thermo Fisher Scientific) or assembled as complementary single-stranded DNA oligos annealed together. Controllers consisting of 8-parts (pA–pH) plus a backbone (pMLC-BB1) were assembled using a standard Golden Gate cloning method (**Figure 2B**) ^27^. Briefly, for each assembly, we started from the 18.5 ng of required part plasmids (pA–pH) and 18.5 ng of the backbone (pMLC-BB1) to be added to a 5 μL Golden Gate reaction. The standard manufacturer’s reaction conditions were used, but at a quarter of their normal volume (New England Biolabs, E1601). 2 μL of this reaction mix was then used to transform 12.5 μL of chemically competent DH10-β cells (New England Biolabs, C3019) for further experiments. All assembled constructs were sequence verified by Sanger sequencing (Eurofins Genomics). Annotated sequences of all part and backbone plasmids and assembled controllers are provided in GenBank format in **Supplementary Data 2**. Plasmid maps are shown in **Supplementary Figures 4** and **5**.

### Characterisation experiments

Single colonies of cells transformed with an appropriate genetic construct were inoculated in 200 μL M9 media supplemented with glucose and kanamycin for selection in a 96-well microtiter plate (Thermo Fisher Scientific, 249952). Cultures were grown for 14 hours in a shaking incubator (Stuart, S1505) at 37 °C and 1250 rpm. Following this, the cultures were diluted 3:40 (15 μL in 185 μL) in M9 media supplemented with glucose, kanamycin for selection and IPTG for induction in a new 96-well microtiter plate and grown for a further 4 hours under the same conditions. Finally, the cultures were further diluted 1:10 (10 μL into 90 μL) in phosphate-buffered saline (PBS) (Gibco,18912-014) containing 2 mg/mL kanamycin to halt protein translation. These samples were incubated at room temperature for 1 hour to allow for full maturation of GFP before flow cytometry was performed.

### Flow cytometry

Measurements of GFP fluorescence in single cells was performed using an Acea Biosciences NovoCyte 3000 flow cytometer equipped with a NovoSampler to allow for automated collection of samples from a 96-well microtiter plate. Data collection was performed using the NovoExpress software. Cells were excited using a 488 nm laser and GFP fluorescence measurements taken using a 530 nm detector. At least 10^6^ events were captured per sample. In addition, to enable conversion of GFP fluorescence into calibrated MEFL units ^58^ a single well per plate contained 15 μL of 8-peak Rainbow Calibration Particles (Spherotech, RCP-30-5A) diluted into 200 μL PBS. Automated gating of events and conversion of GFP fluorescence into MEFL units was performed using the forward (FSC) and side scatter (SSC) channels and the FlowCal Python package version 1.2.2 with default parameters ^58^. To correct for the GFP autofluorescence of cells, *E. coli* DH10-β cells containing no genetic construct were grown in identical conditions. An average measurement of GFP fluorescence in MEFL units from three biological replicates of these cells was then subtracted from fluorescence measurements of cells containing our genetic constructs to correct for cell autofluorescence.

### Plate reader measurements of construct performance in vivo

Single colonies of cells transformed with an appropriate genetic construct were inoculated in 200 μL M9 media supplemented with glucose and kanamycin for selection in a 96-well microtiter plate (Thermo Fisher Scientific, 249952). Cultures were grown for 4 hours in a shaking incubator (Stuart, S1505) at 37 °C and 1250 rpm. Following this, the cultures were diluted 3:40 (15 μL in 185 μL) in M9 media supplemented with glucose, kanamycin for selection and IPTG for induction in a 96-well 190 μm clear base imaging microplate (4titude, Vision Plate^TM^, 4ti-0223). This spectrophotometric assay was performed using a BioTek Synergy Neo2 plate reader at 37°C. Optical density at 600 nm (OD_600_) was measured every 10 min over a 16-hour period. OD_600_ measurements were also taken from samples of M9 medium supplemented with glucose containing no cells to allow for quantification of media autofluorescence. Shaking was automated for the all the length of the experiment. Data collection was performed using the Gen5 version 3.04 software. For each time point, media autofluorescence was subtracted from the sample measurement.

### Cell-free expression

The *E. coli* cell lysate for CFPS was prepared using an autolysis protocol ^59^. In this protocol, *E. coli* BL21-Gold (DE3) cells harboring a pAS-LyseR plasmid give a high-quality cell lysate by freeze-thawing. Specifically, these cells were grown overnight at 37 °C in LB broth supplemented with ampicillin. On the following day, cells were sub-cultured in 2 L of 2X YTPG medium supplemented with ampicillin and grown at 37 °C to an OD_600_ of 1.5. Cells were then harvested at 2000*g* for 15 min at room temperature in four centrifuge bottles and 45 mL of cold S30A buffer (50 mM Tris-HCl at pH 7.7, 60 mM potassium glutamate, 14 mM magnesium glutamate, final pH 7.7) was added to each. Cells were then resuspended by vigorous vortex mixing and poured into a pre-weighted 50 mL falcon tube and centrifuged as in the previous step. The supernatants were completely removed, and the falcon tubes weighted again. The net weight of each pellet was calculated and relative to its weight two volumes of cold S30A supplied with 2 mM DTT was added (3 mL for 1.5 g of pellet). After vigorous vortex mixing, the samples were stored at –80 °C. The next day, frozen cells were placed in a room temperature water bath to thaw, vigorously vortexed, incubated at 37 °C on a shaker for 45 min, vortex mixed again, and then incubated at 37 °C for 45 min. The samples were then centrifuged at 30000*g* for 60 min at 4 °C. The supernatants were carefully pipetted out and aliquoted in 1.5 μL tubes, and then finally centrifuged at 20000*g* using a tabletop centrifuge for 5 min to remove residual cell debris. Aliquots of the lysate were stored at –80 after flash-freezing in liquid nitrogen.

For the prepared lysate, Mg-glutamate and K-glutamate were titrated in with all components of the cell-free reaction based on the protocol of Sun *et al.* ^60^ resulting in concentrations of 10 nM and 60 mM, respectively, for optimal GFP production. Each reaction was prepared at a final volume of 10.5 μL containing 33% lysate, Mg-glutamate and K-glutamate as titrated, and amino acids mix, energy mix, and PEG 8000. For cell-free experiments of the SLC and MLC constructs, maxi-prepped plasmids (using Machery-Nagel NucleoBond Xtra Maxi kit) were added at a final concentration of 10 nM along with varying concentrations of IPTG (0, 0.002, 0.01, 0.05, 0.1, 0.25, 0.5, 1, 2, 5, 10 mM). While gently mixing by pipette, 10 μL of reactions were transferred to a 384-well plate (Greiner Bio-One, 784076) and GFP fluorescence was monitored (excitation/emission wavelengths of 485/528 nM and gain = 100) every 10 min in a plate reader (Tecan Infinite 200 PRO).

### Signal to noise ratio

The signal to noise ratio (SNR) in decibel (dB) units was calculated from the flow cytometry GFP fluorescence distributions using the equation ^23^

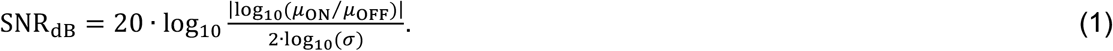

Here, *μ*_ON_ and *μ*_OFF_ are the geometric means of distributions for the ‘on’ and ‘off’ states, respectively, and *σ* is the geometric standard deviation of the distribution for the ‘off’ state. ‘off’ and ‘on’ states correspond to cells grown in 0 and 1 mM IPTG, respectively.

### Data analysis and numerical simulation

Data analysis was performed using Python version 3.7.4 and the NumPy version 1.17.4, SciPy version 1.3.1, Pandas version 1.0.3, FlowCal version 1.2.2, and Matplotlib version 3.1.1 libraries. ODE models were simulated using the odeint function of the SciPy.integrate Python package version 1.1 with default parameters. Steady-state response functions of the controllers were calculated by fitting median GFP fluorescence values from the flow cytometry distributions for a range of input IPTG concentrations to the following Hill function

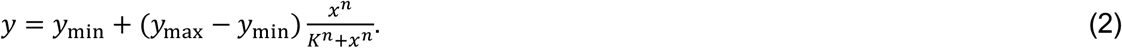

Here, *y* is the output GFP fluorescence in MEFL units, *y*_min_ and *y*_max_ are the minimum and maximum output GFP fluorescence in MEFL units, respectively, *K* is the input IPTG concentration at which the output is half-maximal, *n* is the Hill coefficient, and *x* is the input IPTG concentration. Fitting of the experimental data was performed using non-linear least squares and the curve_fit function from the SciPy.integrate package version 1.1. Genetic diagrams were generated using DNAplotlib version 1.0 ^61,62^ and figures were composed using Omnigraffle version 7.15 and Affinity Designer version 1.8.3.

## Data availability

Python scripts simulating the ODE models of the direct and multi-level controllers be found in **Supplementary Data 1**. Annotated sequences for all plasmids in GenBank format are available in **Supplementary Data 2**. All plasmids used in this study are available from Addgene.

## Acknowledgements

This work was supported by BrisSynBio, a BBSRC/EPSRC Synthetic Biology Research Centre grant BB/L01386X/1 (T.E.G., C.S.G.), a Royal Society PhD Studentship (F.V.G.), a Royal Society University Research Fellowship grant UF160357 (T.E.G.), the Max Planck Society (T.J.E.), and a European Molecular Biology Organization (EMBO) long-term postdoctoral fellowship (A.P.)

## Author Contributions

T.E.G. conceived the study. V.G. designed the genetic toolkit, assembled all controllers and performed the *in vivo* experiments. A.P. performed the *in vitro* cell-free experiments. T.E.G. developed and simulated the mathematical models. V.G. and T.E.G. analysed the data. T.E.G., C.S.G., T.J.E. supervised the work. V.G., T.E.G. and C.S.G. wrote the manuscript with input from all other authors.

