## Supplementary Information for "Harnessing the central dogma for stringent multi-level control of gene expression"

**Supplementary Notes** **Page**

Supplementary Note 1: Derivation of mathematical models 2

Supplementary Note 2: MLC toolkit user guide 3

**Supplementary Figures**

Supplementary Figure 1: Key features of a controller's response function 5

Supplementary Figure 2: Growth curves 6

Supplementary Figure 3: $STAR_2$ performance comparison 7

Supplementary Figure 4: Plasmid maps for the MLC toolkit 8

Supplementary Figure 5: Plasmid maps for the gene expression controllers used 9

**Supplementary Tables**

Supplementary Table 1: Model parameters 10

Supplementary Table 2: Genetic parts used in this study 11

Supplementary Table 3: Performance of controllers in a cell-free expression system 13

**Supplementary References** 14

### Supplementary Note 1: Derivation of mathematical models

To explore how single- and multi-level regulation affected the output protein production rate as a function of an input inducer chemical concentration, we derived mathematical models for each type of system. For the single-level controller we defined equations to track the concentration of GOI transcripts *R* and protein *P* as:

$$\frac{dR}{dt} = \alpha_I - \gamma_R R, \quad (1)$$

$$\frac{dP}{dt} = \alpha_P R - \gamma_P P. \quad (2)$$

Here, $\alpha_I$ is the production rate of GOI transcripts, $\alpha_P$ is the production rate proteins per transcript, and $\gamma_R$ and $\gamma_P$ are first-order degradation rates of the transcripts and output proteins, respectively. In this system, $\alpha_I$ is directly related to the activity of the $P_{L1}$ promoter (**Figure 1C**). To capture a realistic response of a promoter for a small molecule sensor (e.g. the $P_{tac}$ system) that we might use as input to the controller, we made use of empirically measured steady-state response functions that fit to a Hill equation such that

$$\alpha_I = f(x) = y_{\min} + (y_{\max} - y_{\min})\frac{x^n}{K^n + x^n}. \quad (3)$$

Here, *x* is the concentration of the inducer (e.g. a small molecule), $y_{\min}$ and $y_{\max}$ are the minimum and maximum activities of the $P_{L1}$ promoter, *K* is the inducer concentration where the $P_{L1}$ promoter activity is half its maximum, and *n* is the cooperativity.

For the MLC, we extended this model to include additional states for the concentrations of the *L2* regulator *S*, and the complex *C* of the GOI transcript with the *L2* regulator that forms through non-cooperative RNA-RNA hybridisation. By assuming that only these complexes can be translated into output protein, we end up with the following set of equations:

$$\frac{dR}{dt} = \alpha_I - k_{C+}RS + k_{C-}C - \gamma_R R, \quad (4)$$

$$\frac{dS}{dt} = \alpha_I - k_{C+}RS + k_{C-}C - \gamma_S S, \quad (5)$$

$$\frac{dC}{dt} = k_{C+}RS - k_{C-}C - \gamma_C C, \quad (6)$$

$$\frac{dP}{dt} = \alpha_P C - \gamma_P P, \quad (7)$$

where $k_{C+}$ and $k_{C-}$ are the binding and unbinding rates of the GOI transcript with the *L2* RNA-based regulator.

To analyse the behaviour of these systems, numerical simulation of these ODEs was performed using the odeint function of the SciPy.integrate Python package and biologically realistic parameters assuming the input sensor behaved similarly to the $P_{tac}$ system (**Supplementary Table 1**).

### Supplementary Note 2: MLC toolkit user guide

The MLC toolkit is composed of a set of plasmids and assembly rules to create controllers in a combinatorial manner. It includes a backbone plasmid called pMLC-BB1 that is used to hold newly build controllers and a set of 8 types of part plasmid (pA–pH) corresponding to the individual blocks that make up a controller (**Figure 2A**). pMLC-BB1 is based on the pET28a(+) plasmid and contains a pBR322 origin of replication, kanamycin resistance marker and expression cassette for an orange fluorescent protein (*ofp*) gene that is highly visible in transformed cells. The *ofp* expression cassette is flanked by BsaI sites such that it is removed before insertion of a fully assembled MLC. To minimise controller malfunctions due to individual part failure, the orientation of each transcriptional unit has been chosen such that potential transcriptional read-through at any terminator does not cause unwanted expression of other controller elements. The toolkit has also been designed to ensure the accurate and rapid assembly of new designs by minimising the chance of incorrect assembly through the selection of 4 bp overhangs with low cross reactivity and the use of a fluorescent protein drop out upon successful insertion of a controller construct into the backbone to reduce the subsequent screening/sequencing of transformants needed to verify a perfect assembly.

#### *Creating new parts*

Parts for each block of a controller can be created in the form of pre-cloned plasmids, amplicons or even annealed complementary oligos if the length is short. Pre-cloned plasmids are the preferred method as this reduces the chance of part mutations. Amplicons are easy to generate by PCR and allow for the easy addition of 5' and 3' sequences needed for assembly, however, also tend to see higher numbers of mutations. Complementary oligos annealed to create a double stranded DNA part are ideal for smaller sequences up to 60 nt, although again using this method will lead to the highest error rates in the final parts produced. Every new part must be flanked by a standardised sequence that is used to create the necessary single stranded overhangs that are used for correct assembly. These sequences contain a BsaI restriction site and a specific 4 bp overhang. The following table provides details of the necessary 5' and 3' sequences that are needed.

| Part block | Sequence to add at 5' end | Sequence to add at 3' end |
|---|---|---|
| A | NNNNNNGGTCTCA**gtga** | **ggag**TGAGACCNNNNNN |
| B | NNNNNNGGTCTCA**ggag** | **aggt**TGAGACCNNNNNN |
| C | NNNNNNGGTCTCA**aggt** | **gctt**TGAGACCNNNNNN |
| D | NNNNNNGGTCTCA**gctt** | **cgct**TGAGACCNNNNNN |
| E | NNNNNNGGTCTCA**cgct** | **ccgc**CGAGACCNNNNNN |
| F | NNNNNNGGTCTCA**ccgc** | **catc**CGAGACCNNNNNN |

| | | |
|---|---|---|
| G | NNNNNNGGTCTCA**catc** | **tact**TGAGACCNNNNNN |
| H | NNNNNNGGTCTCA**tact** | **tgcc**TGAGACCNNNNNN |

Underlined sequences denote BsaI recognition sites used for Golden Gate assembly. Bold lowercase sequences denote 4 bp overhangs generated after digestion by BsaI which are used for assembly. The six undefined bases at the 5' and 3' ends are optional but help improve digestion efficiency.

***Assembling new controllers***

Assembly of a controller is performed using a standard Golden Gate reaction. We recommend the use of the NEB Golden Gate Assembly Kit (BsaI-HFv2, E1601) and have found that assemblies can be scaled down using this kit to a quarter of their normal volume (5 µL) with still large numbers of successful transformants produced for chemically competent *E. coli* DH10-β cells using 2 µl of the reaction mix. Each reaction should include the backbone plasmid and a single plasmid type for each of the blocks in the controller (**Figure 2A**). Although we rarely see mutations in the part sequences, there is a tendency for mutations to occur within the 4 bp overhangs used for assembly. Therefore, it is recommended that all controllers are fully sequence verified before use.

***Assembling single-level controllers using the MLC toolkit***

Although the toolkit is tailored to produce MLCs, it is possible to create SLCs by substituting several blocks with a specially designed spacer. Specifically, blocks B and C can be replaced by a single spacer element ('BC-spacer', **Supplementary Table 2**) that is able to assemble with block A at its 5'-end and block D at its 3'-end. As mutations in this part are less problematic, we have used annealed oligos to produce this part for creating the $P_{tac}$ SLC. In addition, the removal of the L2 regulator also requires the use of a standard RBS part for block G. We include a strong RBS part in the toolkit for this use ('RBS', **Supplementary Table 2**).

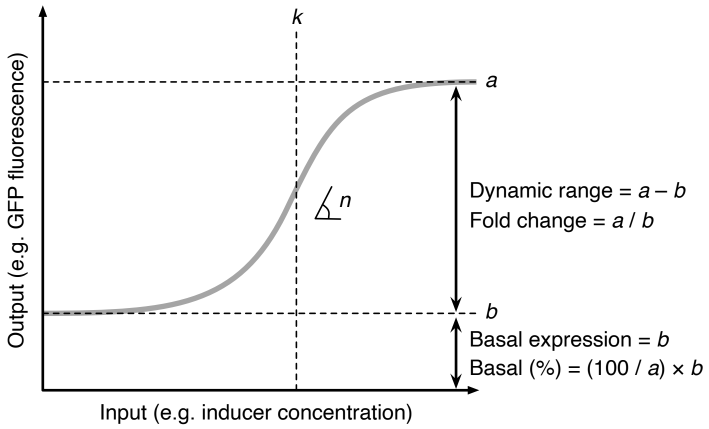


**Supplementary Figure 1: Key features of a controller's response function.** Typical response function for an inducible system shown by thick grey line. The parameters *a*, *b*, *k*, and *n* can be found by fitting to experimental data that measures the controller output across a range of inputs *x* to a Hill function of the form $f(x) = b + (a - b)\left[\frac{x^n}{k^n + x^n}\right]$. Parameter *a* corresponds to the maximum output, *b* to the basal output when no input is present, *k* is the input where half the maximal output minus basal output is achieved, and *n* is the cooperativity (that is manifested as the steepness of the response function).

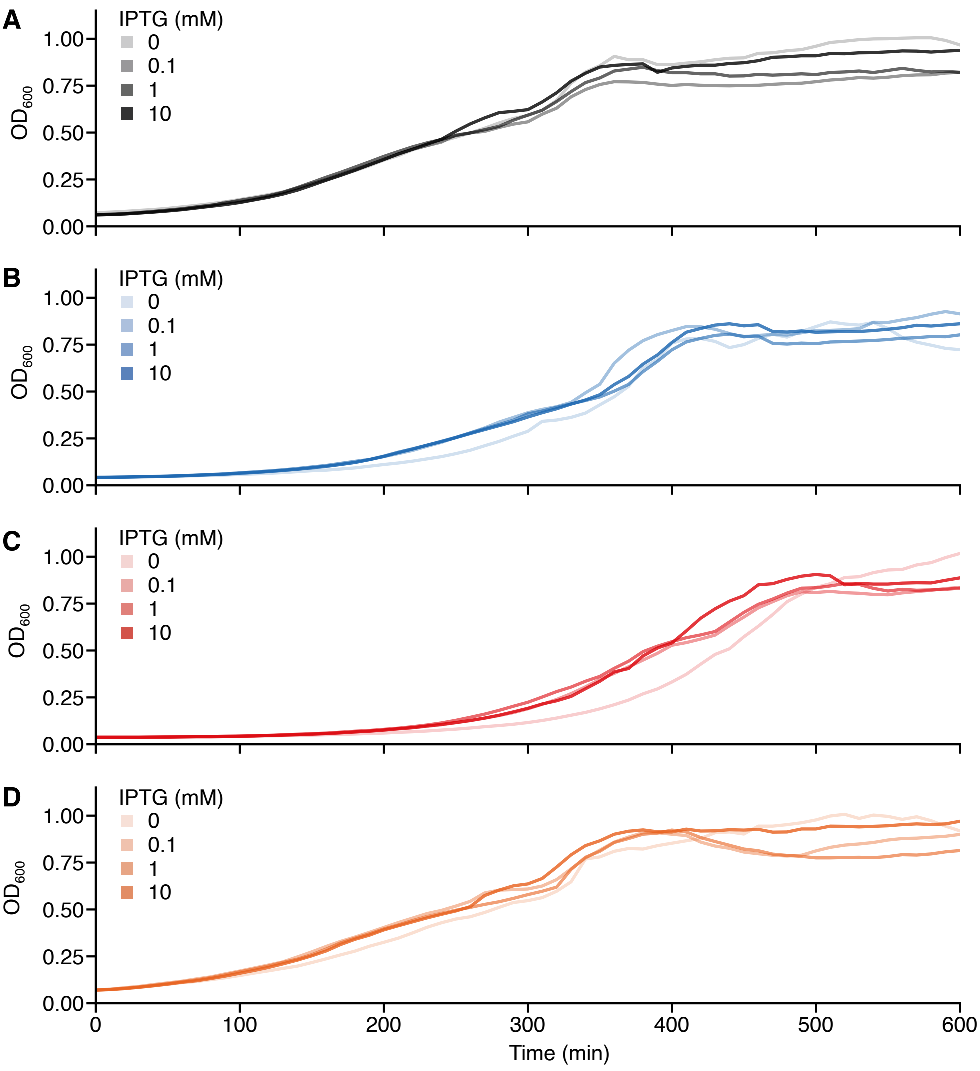


**Supplementary Figure 2: Growth curves.** Time course measurements of the optical density at 600 nm ($OD_{600}$) of *E. coli* DH10-β cells harboring (**A**) pPtac, (**B**) pTHS, (**C**) pSTAR and (**D**) pDC plasmids in the presence of different concentrations of IPTG.

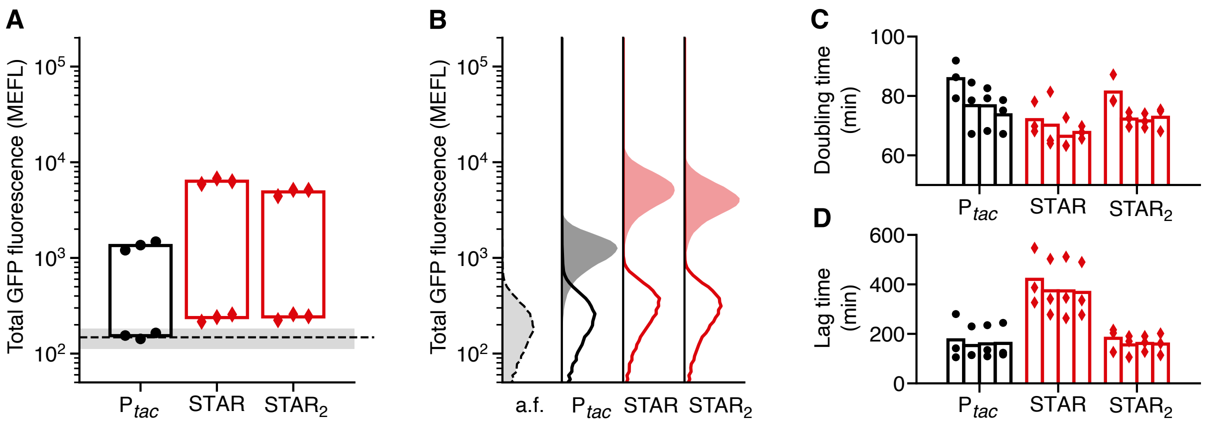


**Supplementary Figure 3: $STAR_2$ performance comparison.** Data for the $P_{tac}$ and STAR designs shown for comparison. (**A**) Total GFP fluorescence for ‘off’ and ‘on’ input states (0 and 1 mM IPTG, respectively). Points show the three biological replicates for each controller and condition (black circles, $P_{tac}$; red diamonds, STAR, $STAR_2$). Black dashed line denotes the mean fluorescence of cell autofluorescence (a.f.) controls containing no plasmid with grey shared region showing ± 1 standard deviation of 11 biological replicates. Fluorescence given in calibrated molecules of equivalent fluorescein (MEFL) units. (**B**) Flow cytometry distributions of total GFP fluorescence for ‘off’ (line) and ‘on’ (shaded) input states. Cell autofluorescence (a.f.) controls containing no controller are shown by black dashed line and light grey filled distributions. (**C**) Doubling time of cells harbouring single- and multi-level controllers for varying concentrations of IPTG (bars left to right for each design: 0, 0.1, 1, 10 mM IPTG). (**D**) Lag time calculated as the time to reach an $OD_{600}$ = 0.15 after inoculation of cells harbouring controllers for varying concentrations of IPTG (bars left to right for each design: 0, 0.1, 1, 10 mM IPTG).

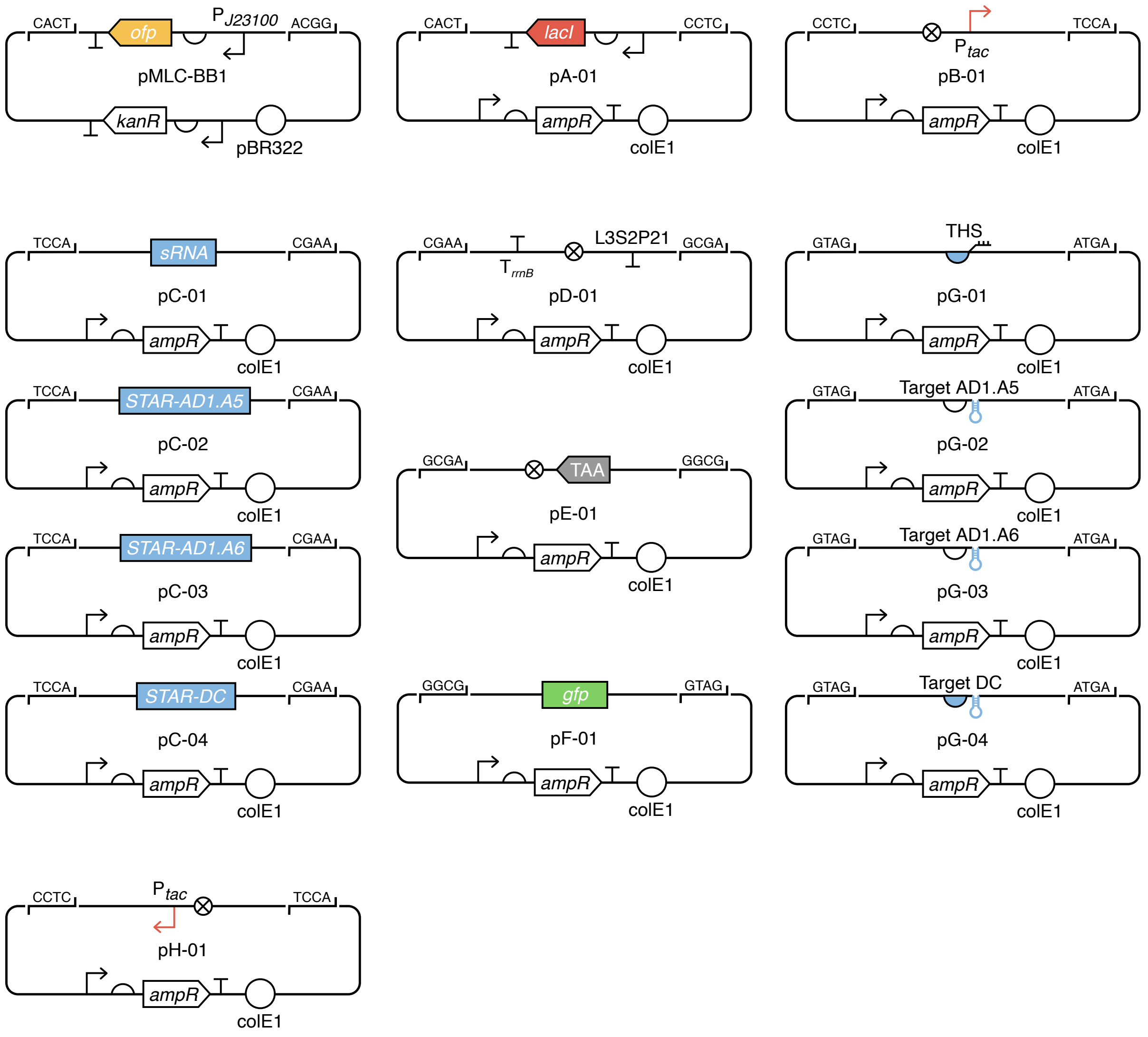


**Supplementary Figure 4: Plasmid maps for the MLC toolkit.** Genetic diagrams are shown using SBOL Visual symbols. BsaI restriction sites flanking each part are shown with their 4 bp overhang sequence after digestion that is used for assembly. The *gfp* part in plasmid pF-01 is in a reverse orientation and lacks a stop codon. The *gfp* open reading frame is completed after assembly with plasmid pE-01 which includes a TAA stop codon.

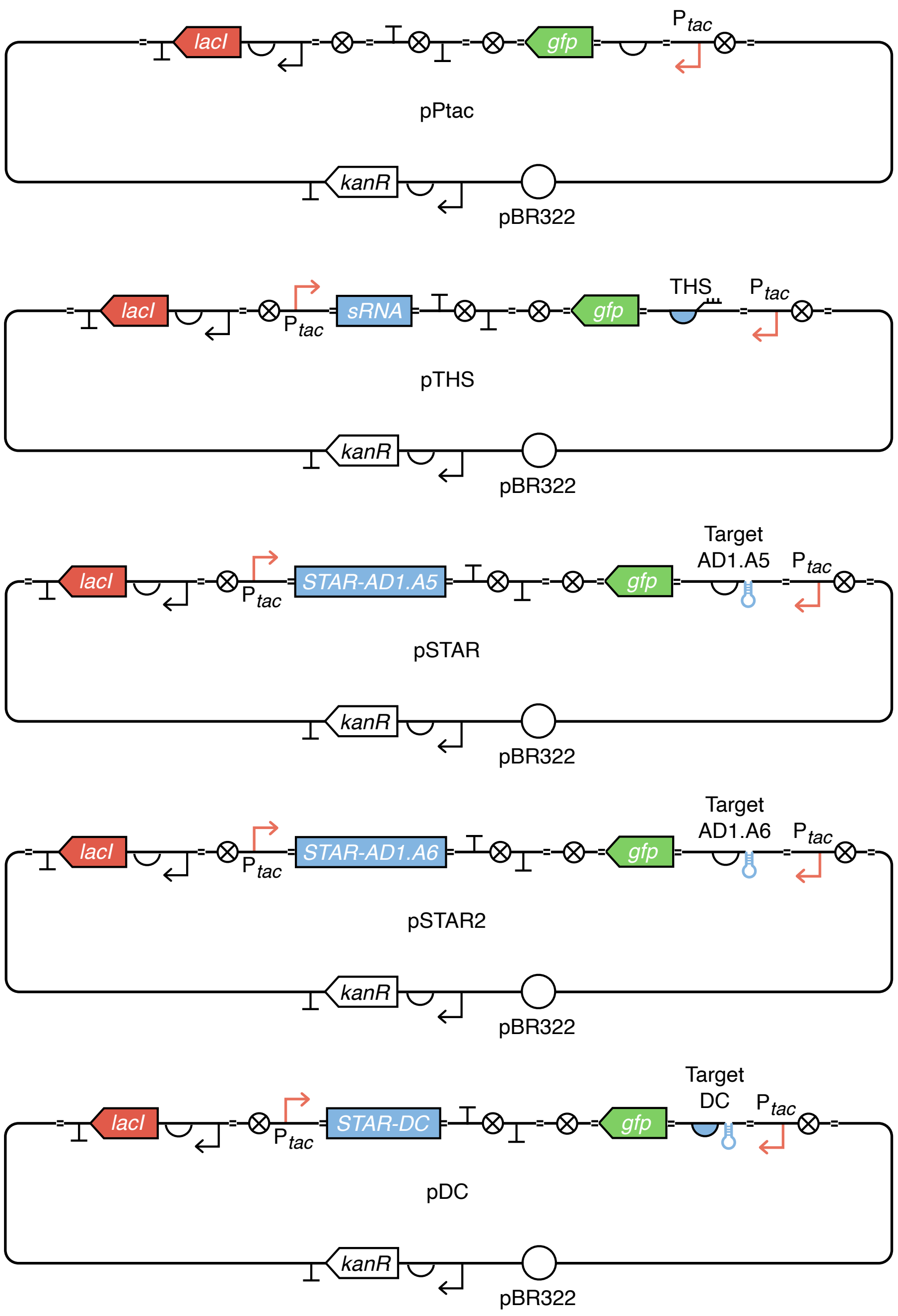


**Supplementary Figure 5: Plasmid maps for the gene expression controllers used.** Genetic diagrams are shown using SBOL Visual symbols and include the scars introduced between core elements of the genetic template.

**Supplementary Table 1: Model parameters**

| Parameter | Description | Value(s) | Units | Refs. |
|---|---|---|---|---|
| $\alpha_I$ | Transcription rate of input promoter | 0.04–100 | transcript min$^{-1}$ | [1,2] |
| $\alpha_P$ | Translation rate of sRNA:transcript complex | 5 | protein complex$^{-1}$ min$^{-1}$ | [3] |
| $k_{C+}$ | sRNA:transcript complex association rate | 0.0257 | complex transcript$^{-1}$ min$^{-1}$ | [4] |
| $k_{C-}$ | sRNA:transcript complex disassociation rate | 0.0067 | transcript complex$^{-1}$ min$^{-1}$ | [4] |
| $\gamma_R$ | Transcript degradation rate | 0.231 | min$^{-1}$ | [5] |
| $\gamma_S$ | sRNA degradation rate | 0.231 | min$^{-1}$ | [5] |
| $\gamma_C$ | sRNA:transcript complex degradation rate | 0.231 | min$^{-1}$ | [5] |
| $\gamma_P$ | Output protein degradation rate | 0.035 | min$^{-1}$ | [6] |
| $y_{\min}$ | Minimum input promoter activity | 0.04 | transcript min$^{-1}$ | [1,2] |
| $y_{\max}$ | Maximum input promoter activity | 100 | transcript min$^{-1}$ | [1,2] |
| $K$ | Concentration of input inducer at which sensor output is half-maximal | 134 | mM | [1] |
| $n$ | Cooperativity of input sensor system | 1.9 | – | [1] |

**Supplementary Table 2: Genetic parts used in this study.**

| Plasmid | Part | Sequence[a] | Ref. |
|---|---|---|---|
| pA1 | *lacI*-TU | GGTCTCA**gtga**AGTGAAAAGAAAAAAGGCCGCAGAGCGGCCTTTTTAGTTAGATCGGAC<br>CCCACATCTCACTGCCCGCTTTCCAGTCGGGAAACCTGTCGTGCCAGCTGCATTAATGA<br>ATCGGCCAACGCGCGGGGAGAGGCGGTTTGCGTATTGGGCGCCAGGGTGGTTTTTCTTT<br>TCACCAGTGAGACTGGCAACAGCTGATTGCCCTTCACCGCCTGGCCCTGAGAGAGTTGC<br>AGCAAGCGGTCCACGCTGGTTTGCCCCAGCAGGCGAAAATCCTGTTTGATGGTGGTTAA<br>CGGCGGGATATAACATGAGCTATCTTCGGTATCGTCGTATCCCACTACCGAGATATCCG<br>CACCAACGCGCAGCCCGGACTCGGTAATGGCGCGCATTGCGCCCAGCGCCATCTGATCG<br>TTGGCAACCAGCATCGCAGTGGGAACGATGCCCTCATTCAGCATTTGCATGGTTTGTTG<br>AAAACCGGACATGGCACTCCAGTCGCCTTCCCGTTCCGCTATCGGCTGAATTTGATTGC<br>GAGTGAGATATTTATGCCAGCCAGCCAGACGCAGACGCGCCGAGACAGAACTTAATGGG<br>CCCGCTAACAGCGCGATTTGCTGGTGACCCAATGCGACCAGATGCTCCACGCCCAGTCG<br>CGTACCGTCCTCATGGGAGTAAATAATACTGTTGATGGGTGTCTGGTCAGAGACATCAA<br>GAAATAACGCCGGAACATTAGTGCAGGCAGCTTCCACAGCAATGGCATCCTGGTCATCC<br>AGCGGATAGTTAATGATCAGCCCACTGACGCGTTGCGCGAGAAGATTGTGCACCGCCGC<br>TTTACAGGCTTCGACGCCGCTTCGTTCTACCATCGACACCACCACGCTGGCACCCAGTT<br>GATCGGCGCGAGATTTAATCGCCGCGACAATTTGCGACGGCGCGTGCAGGGCCAGACTG<br>GAGGTGGCAACGCCAATCAGCAACGACTGTTTGCCCGCCAGTTGTTGTGCCACGCGGTT<br>GGGAATGTAATTCAGCTCCACCATCGCCGCTTCCACTTTTTCCCGCGTTTTCGCAGAAA<br>CGTGGCTGGCCTGGTTCACCACGCGGGAAACGGTCATATAAGAGACACCGGCATACTCT<br>GCGACATCGTATAACGTTACTGGTTTCATATTCACCTCCATGAATTGACTCTCTTCCGG<br>GCGCTATCATGCCATACCGCGAAAGGTTTTTCACCATTCGATGGCGCGCCGCGGCTGGG<br>AATCTGAGACATGAGTCAAGATATTTGCTCGGTAACGTATGCT**ggag**TGAGACC | [7] |
| pB1 | $P_{tac}$ | GGTCTCA**ggag**TGTTGACAATTAATCATCGGCTCGTATAATGTGTGGAATTGTGAGCGC<br>TCACAATT**aggt**TGAGACC | [7] |
| pC1 | STAR-pT1810-DC | GGTCTCA**aggt**AACAAAATAAAGCAATAAGGAATCGCTCACCCAAAGGATCT**gctt**TGA<br>GACC | [8] |
| pC2 | STAR-AD1.A5 | GGTCTCA**aggt**TGAACTGTATACATTCCCCGCTGCTCCAACATTTATACAACTAATTAA<br>AACAATTCACTGTAAAAACT**gctt**TGAGACC | [9] |
| pC3 | STAR-AD1.A6 | GGTCTCA**aggt**TGAACTGTATACATTCCCCGCTGCTCCAACATTTATACAACTAATTAA<br>AACAATTCACTGTAAAAACTTTTCCTAGAC**gctt**TGAGACC | [9] |
| pC4 | sRNA-THS | GGTCTCA**aggt**GGGACCTATTGGACCCGTTTCCAATAGGTGAACAAGACGATAGAACAA<br>GCATTTGCACTTATAGA**gctt**TGAGACC | [10] |
| pD1 | TrrnB-L3S2P21 | GGTCTCA**gctt**ATAAAACGAAAGGCTCAGTCGAAAGACTGGGCCTTTCGTTTTATCTGT<br>TGTTTGTCGGTGAACGCTCTCCTGAGTAGGACAAATCCGCCGGGAGCGGATTTGAACGT<br>TGCGAAGCAACGGCCCGGAGGGTGGCGGGCAGGACGCCCGCCATAAACTGCCAGGCATC<br>AAATTAAGCAGAAGGCCATCCTGACGGATGGCCTTTTTGCCCCCGACCCTTAGACTCTG<br>TACTCAGGGACCAAAACGAAAAAAGGCCCCCCTTTCGGGAGGCCTCTTTTCTGGAATTT<br>GGTACCGAG**cgct**TGAGACC | [11] |
| pE1 | Spacer-TAA | GGTCTCA**cgct**AGTTGTATCGTACGTCGGTCTAGGAATCGTTTAG**ccgc**CGAGACC | – |
| pF1 | *gfp* | GGTCTCA**ccgc**CTCACTTATACAACTCATCCATGCCTAAAGTAATCCCCGCCGCGGTTA<br>CGAACTCCAAAAGGACCATGTGATCACGTTTCTCATTTGGGTCCTTCGATAATTTGGAT<br>TGTGTACTCAGGTAGTGATTGTCAGGAAGTAAAACCGGGCCGTCGCCAATGGGTGTGTT<br>TTGTTGATAATGGTCAGCCAATTGCACGCTGCCATCCTCAATGTTGTGACGAATCTTGA<br>AATTTACCTTAATCCCGTTCTTCTGCTTGTCAGCCATGATATACACGTTGTGCGAATTA<br>TAGTTATACTCCAGTTTATGTCCCAAAATGTTGCCATCTTCTTTAAAGTCAATGCCTTT<br>CAATTCAATGCGATTCACTAACGTATCCCCCTCAAACTTGACCTCGGCGCGTGTCTTAT<br>AGTTCCCGTCATCCTTGAAGAAGATTGTGCGCTCTTGCACATAGCCTTCTGGCATTGCA<br>GATTTAAAGAAGTCGTGCTGCTTCATGTGATCTGGGTAACGCGAGAAACATTGGACCCC<br>GTATGTAAGTGTTGTAACCAGAGTTGGCCAGGGGACTGGAAGTTTTCCTGTAGTACAAA<br>TGAACTTAAGCGTCAATTTGCCGTAAGTAGCGTCGCCTTCGCCCTCGCCGCTAACCGAA<br>AACTTATGACCATTAACGTCCCCATCCAACTCAACCAAGATGGGTACAACGCCTGTAAA<br>CAGCTCCTCTCCTTTACT**catc**TGAGACC | – |
| pG1 | Target-pT181-DC+RBS | GGTCTCA**catc**AAAAAATCGACTCCTTAATCTCAATTTCGTTTAAGGAATCGCTCACCC<br>A**tact**TGAGACC | [8] |
| pG2 | Target-AD1.S5+RBS | GGTCTCA**catc**AGATCCTTCCTCCTAGATCCAAAAAAAAAGCGGGGAATATATACATGA<br>ACTGTATACATTCCCCGCTGCTCCAACATTTATACAACTAATTAAAACAATTCACTGTA<br>AAAACT**tact**TGAGACC | [9] |

| | | | |
|---|---|---|---|
| pG3 | Target-AD1.S6+RBS | GGTCTCA**catc**AGATCCTTCCTCCTAGATCCAAAAAAAAAGCGGGGAATATATACATGAACTGTATACATTCCCCGCTGCTCCAACATTTATACAACTAATTAAAACAATTCACTGTAAAAACTTTTCCTAGAC**tact**TGAGACC | [9] |
| pG4 | Target-THS | GGTCTCA**catc**ACGCTTGTTCTCATGTCTTGTCTCCTCTGTTTCAAGACGATAGAACAAGCATTTGCACTTAT**tact**TGAGACC | [10] |
| pH1 | $P_{tac}$ | GGTCTCA**tact**AATTGTGAGCGCTCACAATTCCACACATTATACGAGCCGATGATTAATTGTCAACA**tgcc**TGAGACC | [7] |
| –[b] | RBS | Fwd: **catc**AGATCCTTCCTCCTAGATCCGCATCCGGGC<br>Rev: **agta**GCCCGGATGCGGATCTAGGAGGAAGGATCT | – |
| –[b] | BC-spacer[c] | Fwd: **ggag**CTCGTTCATCCCGTGGGACATCAAGCTTCGCCTTGATAAA<br>Rev: **aagc**TTTATCAAGGCGAAGCTTGATGTCCCACGGGATGAACGAG | – |

a. Underlined sequences denote BsaI recognition sites used for Golden Gate assembly. Bold lowercase sequences denote 4 bp overhangs generated after digestion by BsaI which are used for assembly.

b. No plasmid was used for these parts. Instead, due to their short length complementary oligos were annealed (**Methods**) to generate double-stranded DNA parts with necessary single-stranded overhangs for assembly.

c. This spacer is used to replace block B and C in an assembly when no *L2* regulation is present.

**Supplementary Table 3: Performance of controllers in a cell-free expression system**[a]

| Controller | Type[b] | Basal[c] (%) | Dynamic range[d] (au/hr) | Fold change[d] | Cooperativity[e], *n* |
|---|---|---|---|---|---|
| $P_{tac}$ | SLC | 10.0 | 192 | 10 | 3.4 |
| THS | MLC | 0.3 | 4481 | 349 | 4.0 |
| STAR | MLC | 1.8 | 855 | 55 | 6.3 |
| DC | MLC | 3.7 | 462 | 27 | 4.1 |

a. All values are averages calculated from three biological replicates. Key performance features of the controllers are visually shown in **Supplementary Figure 1**.

b. SLC refers to 'single-level controller' and MLC refers to 'multi-level controller'.

c. Relative basal expression rate calculated when no IPTG is present and as a percentage of the GFP expression rate for the 'on' state (10 mM IPTG).

d. Calculated between 'on' and 'off' states for cells grown in 0 and 10 mM IPTG, respectively, and given in GFP fluorescence production in arbitrary units per hour (au/hr).

e. From the Hill function fitting of the response functions (**Figure 5A**).
