## Supplementary Data 1 for "Harnessing the central dogma for stringent multi-level control of gene expression": dynamic_model.py

```
#!/usr/bin/env python
"""
Multi-level controller model
============================
    Script that assesses the response of single- and multi-level controllers
    to different temporal inputs.
"""

import numpy as np
import matplotlib
import matplotlib.gridspec as gridspec
import matplotlib.pyplot as plt
from scipy.integrate import odeint

__author__  = 'Thomas E. Gorochowski <>, Biocompute Lab'
__license__ = 'MIT'
__version__ = '1.0'

#############################################################
# SETTINGS/PARAMETERS FOR HOW THE GRAPH LOOKS
#############################################################

# Axes titles
matplotlib.rcParams['axes.labelsize']  = 8
# Numbers on each axis
matplotlib.rcParams['ytick.labelsize']  = 8
matplotlib.rcParams['xtick.labelsize']  = 8
# Space between axis and number
matplotlib.rcParams['ytick.major.pad']  = 0.8
matplotlib.rcParams['ytick.minor.pad']  = 0.8
matplotlib.rcParams['xtick.major.pad']  = 1.5
matplotlib.rcParams['xtick.minor.pad']  = 1.5
matplotlib.rcParams['ytick.direction']  = 'out'
matplotlib.rcParams['xtick.direction']  = 'out'

###############################################################################
# MODEL PARAMETERS (all rates are per min)
###############################################################################

# Dictionary of all the parameters
params_dict = {}
params_dict['a_l2_1_fn'] = lambda t : 0.0
params_dict['a_L2_2_fn'] = lambda t : 0.0
params_dict['a_P'] = 5.0
params_dict['k_f'] = 0.0257
params_dict['k_r'] = 0.00672
params_dict['d_R'] = 0.231
params_dict['d_L2'] = 0.231
params_dict['d_C'] = 0.231
params_dict['d_P'] = 0.035
protein_decay = params_dict['d_P']

###############################################################################
# MODEL DEFINITION
###############################################################################

def single_level_model(y, t, params):
	"""ODE single-level model.
	"""
	# Set interal variables (use readable names)
	a_L2_1_fn = params[0]
	a_P = params[1]
	d_R = params[2]
	d_P = params[3]

	# Set internal states to named variables
	R  = y[0]
	P  = y[1]

	# Calculate the deltas
	dR = a_L2_1_fn(t) - (d_R * R)
	dP = (a_P * R) - (d_P * P)

	# Return the new states
	return [dR, dP]

def make_single_level_params (params_dict):
	"""Create a parameters vector for model from dictionary.
	"""
	params = (
		params_dict['a_L2_1_fn'],
		params_dict['a_P'],
		params_dict['d_R'],
		params_dict['d_P'])
	return params


def multi_level_model(y, t, params):
	"""ODE multi-level model.
	"""
	# Set interal variables (use readable names)
	a_L2_1_fn = params[0]
	a_L2_2_fn = params[1]
	a_P = params[2]
	k_f = params[3]
	k_r = params[4]
	d_R = params[5]
	d_L2 = params[6]
	d_C = params[7]
	d_P = params[8]

	# Set internal states to named variables
	R  = y[0]
	L2 = y[1]
	C  = y[2]
	P  = y[3]

	# Calculate the deltas
	dR = a_L2_1_fn(t) - (k_f * R * L2) + (k_r * C) - (d_R * R)
	dL2 = a_L2_2_fn(t) - (k_f * R * L2) + (k_r * C) - (d_L2 * L2)
	dC = (k_f * R * L2) - (k_r * C) - (d_C * C)
	dP = (a_P * C) - (d_P * P)

	# Return the new states
	return [dR, dL2, dC, dP]

def make_multi_level_params (params_dict):
	"""Create a parameters vector for model from dictionary.
	"""
	params = (
		params_dict['a_L2_1_fn'],
		params_dict['a_L2_2_fn'],
		params_dict['a_P'],
		params_dict['k_f'],
		params_dict['k_r'],
		params_dict['d_R'],
		params_dict['d_L2'],
		params_dict['d_C'],
		params_dict['d_P'])
	return params

###############################################################################
# MODEL SIMULATION
###############################################################################

# Shared parameters
tcrit_points=[100.0, 101, 150.0, 151]
t_start = 80
t_output = np.arange(0.0, 250.0, 1.0)
y0_single = [0.001, 5.5]
y0_multi = [0.001, 0.001, 0.001, 0.025]

fig_x = 1.25
left_per = 0.4
fig_size = (fig_x, 0.7)

y_lim = [-0.3, 800]
linthreshy = 1.0
annotate_lw = 0.8
red_col = (0.95,0.30,0.25)

###############################################################################

def run_single_level_sim (a_L2_input_fn, model, t_output, y0):
	params_dict['a_L2_1_fn'] = a_L2_input_fn
	# Create an array to hold the parameters
	params = make_single_level_params(params_dict)
	# Copy the parameters for each run
	new_params = list(params)
	y0_copy = list(y0)
	y_out = odeint(model, y0_copy, t_output, args=(new_params,), tcrit=tcrit_points)
	return y_out

def run_multi_level_sim (a_L2_1_input_fn, a_L2_2_input_fn, model, t_output, y0):
	params_dict['a_L2_1_fn'] = a_L2_1_input_fn
	params_dict['a_L2_2_fn'] = a_L2_2_input_fn
	# Create an array to hold the parameters
	params = make_multi_level_params(params_dict)
	# Copy the parameters for each run
	new_params = list(params)
	y0_copy = list(y0)
	y_out = odeint(model, y0_copy, t_output, args=(new_params,), tcrit=tcrit_points)
	return y_out

###############################################################################

def input_pulse_fn (t):
	if t >= 100.0 and t < 101:
		return 2.0
	if t >= 150.0 and t < 151:
		return 2.0
	return 0.01

y_out_single = run_single_level_sim (input_pulse_fn, single_level_model, t_output, y0_single)
y_out_multi = run_multi_level_sim (input_pulse_fn, input_pulse_fn, multi_level_model, t_output, y0_multi)

fig = plt.figure(figsize=fig_size)
gs = gridspec.GridSpec(1, 1)
ax = plt.subplot(gs[0])
ax.plot(t_output[t_start:], np.array(y_out_single[t_start:, -1])*protein_decay, color=(0.0,0.0,0.0), lw=1.2, zorder=-2)
ax.plot(t_output[t_start:], np.array(y_out_multi[t_start:, -1])*protein_decay, color=(0.6,0.6,0.6), lw=1.2, zorder=-1)
ax.set_yscale('symlog', linthreshy=linthreshy)
ax.set_ylim(y_lim)
ax.set_xlim([min(t_output[t_start:]), max(t_output-30)])
ax.spines['bottom'].set_visible(False)
ax.axes.get_xaxis().set_visible(False)
ax.spines['top'].set_visible(False)
ax.spines['right'].set_visible(False)
plt.subplots_adjust(hspace=.0 , wspace=.00, left=left_per, right=.95, top=.95, bottom=.25)
fig.savefig('./dynamic_response_pulse.png', dpi=200)
plt.close('all')

###############################################################################

def input_step_fn (t):
	if t > 100:
		return 5.0
	return 0.01

y_out_single = run_single_level_sim (input_step_fn, single_level_model, t_output, y0_single)
y_out_multi = run_multi_level_sim (input_step_fn, input_step_fn, multi_level_model, t_output, y0_multi)

fig = plt.figure(figsize=fig_size)
gs = gridspec.GridSpec(1, 1)
ax = plt.subplot(gs[0])
ax.plot(t_output[t_start:], np.array(y_out_single[t_start:, -1])*protein_decay, color=(0.0,0.0,0.0), lw=1.2, zorder=-2)
ax.plot(t_output[t_start:], np.array(y_out_multi[t_start:, -1])*protein_decay, color=(0.6,0.6,0.6), lw=1.2, zorder=-1)
#ax.axhline(y_out_single[t_start, -1]*protein_decay, linewidth=annotate_lw, linestyle='--', color=red_col, zorder=-100)
ax.set_yscale('symlog', linthreshy=linthreshy)
ax.set_ylim(y_lim)
ax.set_xlim([min(t_output[t_start:]), max(t_output-30)])
ax.spines['bottom'].set_visible(False)
ax.axes.get_xaxis().set_visible(False)
ax.spines['top'].set_visible(False)
ax.spines['right'].set_visible(False)
ax.tick_params(labelleft=False)
plt.subplots_adjust(hspace=.0 , wspace=.00, left=left_per, right=.95, top=.95, bottom=.25)
fig.savefig('./dynamic_response_step.png', dpi=200)
plt.close('all')

###############################################################################

def input_square_fn (t):
	if t > 100.0 and t < 130.0:
		return 5.0
	return 0.01

y_out_single = run_single_level_sim (input_square_fn, single_level_model, t_output, y0_single)
y_out_multi = run_multi_level_sim (input_square_fn, input_square_fn, multi_level_model, t_output, y0_multi)

fig = plt.figure(figsize=fig_size)
gs = gridspec.GridSpec(1, 1)
ax = plt.subplot(gs[0])
ax.plot(t_output[t_start:], np.array(y_out_single[t_start:, -1])*protein_decay, color=(0.0,0.0,0.0), lw=1.2, zorder=-2)
ax.plot(t_output[t_start:], np.array(y_out_multi[t_start:, -1])*protein_decay, color=(0.6,0.6,0.6), lw=1.2, zorder=-1)
#ax.axhline(y_out_single[t_start, -1]*protein_decay, linewidth=annotate_lw, linestyle='--', color=red_col, zorder=-100)
ax.set_yscale('symlog', linthreshy=linthreshy)
ax.set_ylim(y_lim)
ax.set_xlim([min(t_output[t_start:]), max(t_output-30)])
ax.spines['bottom'].set_visible(False)
ax.axes.get_xaxis().set_visible(False)
ax.spines['top'].set_visible(False)
ax.spines['right'].set_visible(False)
ax.tick_params(labelleft=False)
plt.subplots_adjust(hspace=.0 , wspace=.00, left=left_per, right=.95, top=.95, bottom=.25)
fig.savefig('./dynamic_response_square.png', dpi=200)
plt.close('all')

###############################################################################
# Plot all inputs

fig_size_input = (fig_x, 1.0)
#fig_size_input = (2.2, 0.5)

out = list(map(lambda x: [input_square_fn(x)], t_output[t_start:]))
fig = plt.figure(figsize=fig_size_input)
gs = gridspec.GridSpec(1, 1)
ax = plt.subplot(gs[0])
ax.plot(t_output[t_start:], out, color=(0.0,0.0,0.0), lw=1.2, zorder=-2)
ax.set_ylim([-2, 10])
ax.set_xlim([min(t_output[t_start:]), max(t_output-30)])
ax.tick_params(labelleft=False)
ax.spines['top'].set_visible(False)
ax.spines['right'].set_visible(False)
ax.set_yticks([0, 5, 10])
ax.set_yticklabels(['0', '5', '10'])
plt.subplots_adjust(hspace=.0 , wspace=.00, left=left_per, right=.95, top=.9, bottom=.41)
fig.savefig('./dynamic_input_square.png', dpi=200)
plt.close('all')

out = list(map(lambda x: [input_step_fn(x)], t_output[t_start:]))
fig = plt.figure(figsize=fig_size_input)
gs = gridspec.GridSpec(1, 1)
ax = plt.subplot(gs[0])
ax.plot(t_output[t_start:], out, color=(0.0,0.0,0.0), lw=1.2, zorder=-2)
ax.set_ylim([-2, 10])
ax.set_xlim([min(t_output[t_start:]), max(t_output-30)])
ax.tick_params(labelleft=False)
ax.spines['top'].set_visible(False)
ax.spines['right'].set_visible(False)
ax.set_yticks([0, 5, 10])
ax.set_yticklabels(['0', '5', '10'])
plt.subplots_adjust(hspace=.0 , wspace=.00, left=left_per, right=.95, top=.9, bottom=.41)
fig.savefig('./dynamic_input_step.png', dpi=200)
plt.close('all')

out = list(map(lambda x: [input_pulse_fn(x)], t_output[t_start:]))
fig = plt.figure(figsize=fig_size_input)
gs = gridspec.GridSpec(1, 1)
ax = plt.subplot(gs[0])
ax.plot(t_output[t_start:], out, color=(0.0,0.0,0.0), lw=1.2, zorder=-2)
ax.set_ylim([-2, 10])
ax.set_xlim([min(t_output[t_start:]), max(t_output-30)])
ax.spines['top'].set_visible(False)
ax.spines['right'].set_visible(False)
ax.set_yticks([0, 5, 10])
ax.set_yticklabels(['0', '5', '10'])
plt.subplots_adjust(hspace=.0 , wspace=.00, left=left_per, right=.95, top=.9, bottom=.41)
fig.savefig('./dynamic_input_pulse.png', dpi=200)
plt.close('all')
```
