## Supplementary Data 1 for "Harnessing the central dogma for stringent multi-level control of gene expression": ss_response_model.py

```
#!/usr/bin/env python
"""
Multi-level controller model
============================
    Script that generates a steady state response function for the 
    single- and multi-level controllers.
"""

import numpy as np
import matplotlib
import matplotlib.gridspec as gridspec
import matplotlib.pyplot as plt
from scipy.integrate import odeint

__author__  = 'Thomas E. Gorochowski <>, Biocompute Lab'
__license__ = 'MIT'
__version__ = '1.0'

#############################################################
# SETTINGS/PARAMETERS FOR HOW THE GRAPH LOOKS
#############################################################

# Axes titles
matplotlib.rcParams['axes.labelsize']  = 8
# Numbers on each axis
matplotlib.rcParams['ytick.labelsize']  = 8
matplotlib.rcParams['xtick.labelsize']  = 8
# Space between axis and number
matplotlib.rcParams['ytick.major.pad']  = 0.8
matplotlib.rcParams['ytick.minor.pad']  = 0.8
matplotlib.rcParams['xtick.major.pad']  = 1.5
matplotlib.rcParams['xtick.minor.pad']  = 1.5
matplotlib.rcParams['ytick.direction']  = 'out'
matplotlib.rcParams['xtick.direction']  = 'out'

###############################################################################
# MODEL PARAMETERS (all rates are per min)
###############################################################################

# Dictionary of all the parameters
params_dict = {}
params_dict['a_l2_1_fn'] = lambda t : 0.0
params_dict['a_L2_2_fn'] = lambda t : 0.0
params_dict['a_P'] = 5.0
params_dict['k_f'] = 0.0257
params_dict['k_r'] = 0.00672
params_dict['d_R'] = 0.231
params_dict['d_L2'] = 0.231
params_dict['d_C'] = 0.231
params_dict['d_P'] = 0.035
protein_decay = params_dict['d_P']

###############################################################################
# MODEL DEFINITION
###############################################################################

def single_level_model(y, t, params):
	"""ODE single-level model.
	"""
	# Set interal variables (use readable names)
	a_L2_1_fn = params[0]
	a_P = params[1]
	d_R = params[2]
	d_P = params[3]

	# Set internal states to named variables
	R  = y[0]
	P  = y[1]

	# Calculate the deltas
	dR = a_L2_1_fn(t) - (d_R * R)
	dP = (a_P * R) - (d_P * P)

	# Return the new states
	return [dR, dP]

def make_single_level_params (params_dict):
	"""Create a parameters vector for model from dictionary.
	"""
	params = (
		params_dict['a_L2_1_fn'],
		params_dict['a_P'],
		params_dict['d_R'],
		params_dict['d_P'])
	return params


def multi_level_model(y, t, params):
	"""ODE multi-level model.
	"""
	# Set interal variables (use readable names)
	a_L2_1_fn = params[0]
	a_L2_2_fn = params[1]
	a_P = params[2]
	k_f = params[3]
	k_r = params[4]
	d_R = params[5]
	d_L2 = params[6]
	d_C = params[7]
	d_P = params[8]

	# Set internal states to named variables
	R  = y[0]
	L2 = y[1]
	C  = y[2]
	P  = y[3]

	# Calculate the deltas
	dR = a_L2_1_fn(t) - (k_f * R * L2) + (k_r * C) - (d_R * R)
	dL2 = a_L2_2_fn(t) - (k_f * R * L2) + (k_r * C) - (d_L2 * L2)
	dC = (k_f * R * L2) - (k_r * C) - (d_C * C)
	dP = (a_P * C) - (d_P * P)

	# Return the new states
	return [dR, dL2, dC, dP]

def make_multi_level_params (params_dict):
	"""Create a parameters vector for model from dictionary.
	"""
	params = (
		params_dict['a_L2_1_fn'],
		params_dict['a_L2_2_fn'],
		params_dict['a_P'],
		params_dict['k_f'],
		params_dict['k_r'],
		params_dict['d_R'],
		params_dict['d_L2'],
		params_dict['d_C'],
		params_dict['d_P'])
	return params

###############################################################################
# MODEL SIMULATION
###############################################################################

def run_single_level_ss_sim_range (a_L2_range, model):
	y0_base = [0.01, 0.0]
	data = []
	for a_L2 in a_L2_range:
		params_dict['a_L2_1_fn'] = lambda t : a_L2
		# Create an array to hold the parameters
		params = make_single_level_params(params_dict)
		# Copy the parameters for each run
		new_params = list(params)
		t_output = np.arange(0.0, 1000.0, 1.0)
		y0 = list(y0_base)
		y_out = odeint(model, y0, t_output, args=(new_params,))
		data.append( y_out[-1][1] )
	return data

def run_muti_level_ss_sim_range (a_L2_range, model):
	y0_base = [0.01, 0.01, 0.0, 0.0]
	data = []
	for a_L2 in a_L2_range:
		params_dict['a_L2_1_fn'] = lambda t : a_L2
		params_dict['a_L2_2_fn'] = lambda t : a_L2
		# Create an array to hold the parameters
		params = make_multi_level_params(params_dict)
		# Copy the parameters for each run
		new_params = list(params)
		t_output = np.arange(0.0, 1000.0, 1.0)
		y0 = list(y0_base)
		y_out = odeint(model, y0, t_output, args=(new_params,))
		data.append( y_out[-1][3] )
	return data

# Generate realistic P_tac production rate based on IPTG concentration (mM)
def hill_fn (x, y_min, y_max, k, n):
	return y_min + (y_max - y_min)*(np.power(x, n)/(np.power(k, n) + np.power(x, n)))
p_Tac_params = [1.22596888e-03, 2.72767581e+00, 1.34275264e+02, 1.92555987e+00]
fit_x_data = np.logspace(-1.0, 3.3, 200)
fit_y_data = hill_fn(fit_x_data, p_Tac_params[0], p_Tac_params[1], p_Tac_params[2], p_Tac_params[3])
fit_y_data = (fit_y_data/2.72767581)*100.0

# Collect the data
response_single = run_single_level_ss_sim_range (fit_y_data, single_level_model)
response_multi = run_muti_level_ss_sim_range (fit_y_data, multi_level_model)

def simpleaxis(ax):
    ax.spines['top'].set_visible(False)
    ax.spines['right'].set_visible(False)
    ax.get_xaxis().tick_bottom()
    ax.get_yaxis().tick_left()

# Plot the figure
fig = plt.figure(figsize=(1.12, 1.2))
gs = gridspec.GridSpec(1, 1)
ax = plt.subplot(gs[0])
simpleaxis(ax)
# This is the production rate (protein/min)
plt.plot(fit_x_data, np.array(response_single)*protein_decay, color=(0.0,0.0,0.0), lw=1.2, zorder=-2)
plt.plot(fit_x_data, np.array(response_multi)*protein_decay, color=(0.6,0.6,0.6), lw=1.2, zorder=-1)
ax.set_xscale('log')
ax.set_yscale('log')
ax.set_ylim([0.01, 10000])
ax.set_xlim([min(fit_x_data), max(fit_x_data)])
plt.subplots_adjust(hspace=.0 , wspace=.00, left=.25, right=.95, top=.95, bottom=.2)
fig.savefig('./ss_response.png', dpi=200)
plt.close('all')
```
